# Cell group analysis reveals changes in upper-layer neurons associated with schizophrenia

**DOI:** 10.1101/2020.10.22.351213

**Authors:** Rujia Dai, Lulu Chen, Sihan Liu, Chiung-Ting Wu, Yu Chen, Yi Jiang, Jiacheng Dai, Qihang Wang, Richard Kopp, Guoqiang Yu, Yue Wang, Chao Chen, Chunyu Liu

## Abstract

Genome-wide association studies (GWAS) of schizophrenia (SCZ) have revealed over 100 risk loci. We investigated whether these SCZ-associated variants regulate gene expression by cell type. Using a fully unsupervised deconvolution method, we calculated gene expression by clusters of estimated cell types (cell-groups, CGs). Five CGs emerged in the dorsolateral prefrontal cortices (DLPFC) of 341 donors with and without SCZ. By mapping expression quantitative trait loci (eQTL) per CG, we partitioned the heritability of SCZ risk in GWAS by CGs. CG-specific expressions and eQTLs were replicated in both a deconvoluted bulk tissue data set with a different method and also in sorted-cell expression data. Further, we characterized CG-specific gene differential expression and cell proportion changes in SCZ brains. We found upper-layer neurons in the DLPFC to be associated with SCZ based on enrichment of SCZ heritability in eQTLs, disease-related transcriptional signatures, and decreased cell proportion. Our study suggests that neurons and related anomalous circuits in the upper layers of the DLPFC may have a major contribution to SCZ risk.

## Introduction

Schizophrenia (SCZ) is a complex psychiatric disorder, with approximately 80% of phenotypic variation explained by genetics^1^. In recent years, a surge of genetic markers associated with SCZ has flowed from genome-wide association studies (GWAS)^2^. Nonetheless, interpreting the biological meaning of these GWAS signals remains a challenge. SCZ GWAS loci are located largely within non-coding regions of the genome, pointing to a mechanistic involvement in gene regulation^3^. Expression quantitative trait loci (eQTL) mapping provides an effective means for connecting and interpreting GWAS results with underlying relevant expression regulation^4^. The PsychENCODE^5^, CommonMind^6^, Brain Cloud^7^, and UK Brain Expression Consortium^8^ projects produced large-scale eQTL resources. These data have been integrated with GWAS in the effort to pinpoint risk genes^9–11^.

A primary pitfall of current brain eQTL analyses, however, is the indifference to cell-type specificity^12^. All existing brain eQTL resources originate from homogenate bulk tissues in the brain, a mixture of many cell types or subtypes. Bulk tissue eQTL mapping offers only averaged effects across the composite cell types. It is therefore confounded by variations in proportion and gene expression. Reports of cell-type specificity of eQTLs from blood^13^ provide evidence that at least some common genetic variants shape gene expression by cell types^14^. However, the cell-type-specific eQTLs in the human brain have not yet been reported.

To date, the majority of transcriptomic studies of SCZ have similarly relied on data derived from bulk tissue, using differential analysis and co-expression analyses^15,16^. Such studies bear two major limitations: 1) it is unclear whether genes that are differentially expressed between individuals with SCZ and controls are attributed to differences in expression or changes in cell proportion, and 2) it is unknown whether changes in SCZ-related expression occur only within one specific cell type or multiple cell types.

Researchers investigate cell-type-specific expression using either of two approaches: 1) sequencing individual or sorted cells using single-cell or single-nuclei sequencing (scRNA-seq or snRNA-seq), or 2) the computational deconvolution of bulk tissue transcriptomic data. To date, fewer than 50 human brain samples have been sequenced and published using scRNA-seq or snRNA-seq^17,18^. For the time being, such methods are impractical for most genetic or case-control studies due to the large sample size and associated costs required to achieve adequate statistical power.^19^ Alternatively, the computational approach can deconvolute bulk tissue data into cell-type-specific data with sufficient accuracy^20^.

The computational deconvolution of bulk tissue data can estimate cell type proportion and cell type-specific expression in either a supervised or an unsupervised manner. Supervised deconvolution relies on an accurate reference, for example, cell type-specific expression from the same species and brain region. However, suitable references are often unavailable, and whether references developed from control samples can be used to deconvolute patient data is uncertain. In contrast, using unsupervised deconvolution sidesteps this problem by estimating cell-type-specific expression and cell proportion without prior information. Unsupervised deconvolution classifies a group of cells algorithmically to differentiate them from other groups. Since these groups may not always reflect biologically homogeneous cell types, we instead use the term cell groups (CGs). The resulting CGs represent an estimated cluster of cell sub-types extractable from bulk tissue. Using unsupervised deconvolution methods, like convex analysis of mixture (CAM),^21,22^ identifies CGs based on gene expression patterns from all tested individuals.

Some advanced deconvolution methods can *infer* cell-type-specific expression per individual, based upon the methylation^23^ or gene expression^24^ profile from the donor’s bulk tissue. Again, such supervised methods rely on suitable references. Ideally, sample-wise and unsupervised deconvolution methods are needed to investigate CG-specific expressions from bulk tissue data in a cost-effective manner.

We hypothesized that GWAS variants associated with the risk for SCZ regulate gene expression within one or more specific brain cell types. To identify CG-specific expressions at the sample level, we applied the newly-developed method^25,26^ sample-wise convex analysis of mixtures (swCAM) to RNA-seq data from 93 postmortem brain samples of donors with SCZ and 248 healthy donors (Supplemental fig.1). An independent postmortem brain dataset of 605 samples was used to replicate the swCAM deconvolution results. Further, we generated CG-specific eQTLs and tested whether the CG-specific eQTLs were enriched for SCZ GWAS signals. To detect SCZ-related transcriptional changes per CG, we performed differential expression analyses, followed by co-expression network and transcriptome-wide association analyses of the deconvoluted expression data. We integrated results from GWAS enrichment, cell proportion, and transcriptomic changes to target SCZ-associated CGs. The processed data and results are accessible on our website (http://lbpg.upstate.edu/module_search/).

## Results

### Sample-wise deconvolution of RNA-seq data from bulk brain tissue of donors with and without SCZ

Five CGs emerged from swCAM deconvolution of bulk tissue data. Their identities were first annotated by enrichment testing of known cell-type marker genes. We calculated the enrichment of marker genes of major brain cell types^27^ (Supplemental table 1) in the top cell-type differentially expressed genes (ctDEG) for each CG. Top ctDEGs were selected by differential expression analysis of target CG versus the remainder of CGs (log2FC>2, False Discovery Rate [FDR] q <0.05, Wilcoxon signed-rank test). The five annotated CGs represented astrocytes, two neuronal groups, a mixture of astrocytes, microglia, and endothelial cells, and oligodendrocytes (p value <0.05, Fisher’s exact test, Fig. 1A, Supplemental Fig. 2). We also used Expression Weighted Cell Type Enrichment (EWCE) test as a replication for annotating CGs. We applied EWCE to two snRNA-seq data sets^17,28^ from the human frontal cortex and tested to determine if ctDEGs are expressed more highly in a given cell type than that by chance. The EWCE results were consistent with previous CG annotations except for CG1 (Fig. 1B). Because CG1 was enriched for two glial cell subtypes in the EWCE test of two snRNA-seq data sets, we defined CG1 as a mixture of astrocyte and endothelial cells.

**Fig. 1.**
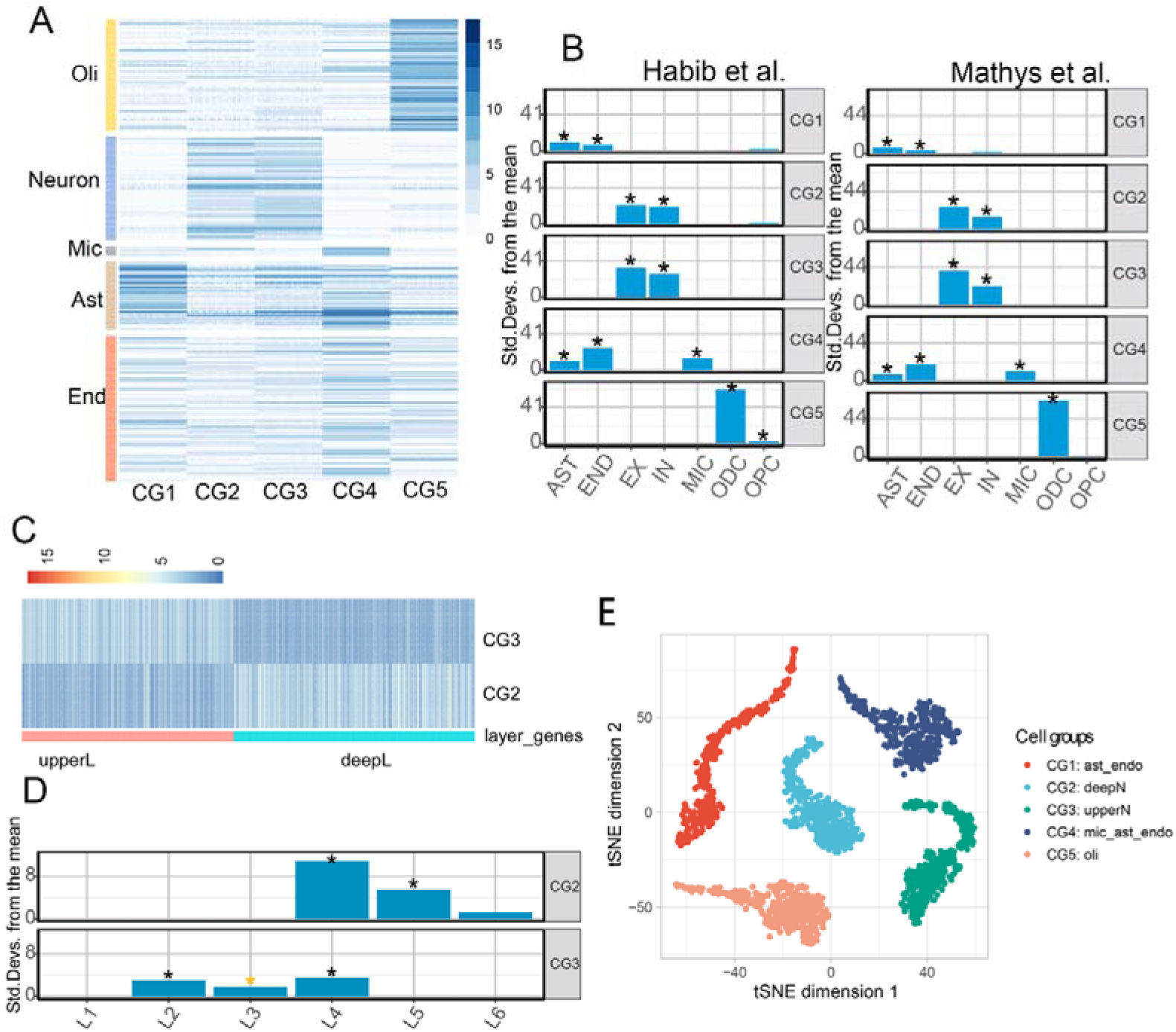
Cell-group-specific expression for each individual. (A) Expression of brain cell-type marker genes per individual estimated cell groups (CGs). Oli: oligodendrocytes, Mic: microglia, Ast: astrocytes, Endo: endothelial cells. (B) Annotation of CGs with Expression Weighted Cell Type Enrichment (EWCE) test. Two snRNA-seq data sets from human frontal cortex were used. The black asterisks denote enrichment p value <0.05. (C) Gradients of gene expression across cortical layers in deconvoluted neuronal CGs. The red column denotes upper-layer genes (upperL genes), which express in a decreasing gradient from layers 1 to 6, and the blue column denotes deep-layer genes (deepL genes), which express in an increasing gradient from layers 1 to 6 (FDR<0.05, Spearman correlation). (D) EWCE results showed neuronal CGs enriched in neurons of different cortical layers. The black asterisks denote enrichment p value <0.05 and the yellow asterisk denotes nominal p value <0.05. (E) tSNE plot of deconvoluted groups, color-coded by the annotated CG. Each dot denotes one deconvoluted cell within a sample. Ast_endo: mixture CG of astrocyte and endothelial cells, deepN: deep-layer neuron CG, upperN: upper-layer neuron CG, mic_ast_endo: mixture CG of microglia, astrocyte, and endothelial cells, oli: oligodendrocyte CG.

To test the expression similarity between estimated CGs and corresponding cell types, we performed a Spearman correlation test between CG expressions and snRNA-seq data from the human frontal cortex^28^. We corrected the batch effects between the tested data prior to the correlation test. The three CGs that only enriched for single cell types showed high correlations with corresponding cell types (averaged rho_neuron_CG2_ =0.83, rho_neuron_CG3_ =0.81, rho_oligodendrocyte_CG5_=0.86, all three p values < 0.05, Supplemental Fig. 3) The averaged correlation coefficient for two glial mixture CGs was 0.55 (SD=0.15).

To further differentiate the two neuronal clusters, marker gene-sets of standard neuronal subtypes were applied. We initially suspected that CG2 and CG3 represented inhibitory and excitatory neurons but were dissuaded because there was not convincing statistical differentiation provided by marker genes of these neuron types (Fisher’s exact test p>0.05, Supplemental Fig. 4). We then speculated that CG2 and CG3 may reflect the origin of specific cortical layers. With only two detected neuronal CGs and six cortical layers, we took a quantitative approach in allocating gene expression to gradient cortical layers: 770 *upper-layer (upperL) genes* were expressed most abundantly in the uppermost layer (layer 1) steadily decreasing by layer with the least expression in the layer 6, while 875 *deep-layer (deepL) genes* were distributed in the opposite direction (FDR<0.05, linear regression). *UpperL* and *deepL genes* corresponded to the CG2 and CG3 clusters respectively (Fisher’s exact test, p_upperL_<2.2e-16, p_deepL_=7.90e-10, Fig. 1C).

The expression data of manually isolated neurons from each of the six layers of the middle temporal cortex^29^ was used to validate the identity of neuronal CGs. We calculated the CG-specific difference for each gene by comparing the expression of CG2 and CG3 individually to that of the other CGs. Layer-specific differences were also calculated for each gene by comparing the neuronal expression of one specific layer to that of the other layers. CG-specific differences and layer-specific differences were then tested for correlation. CG3-specific differences were highly correlated to layer-specific differences of neurons from layers 1 through 3, while CG3-specific differences were poorly correlated to layer-specific differences of neurons from layers 4 through 6. In contrast, correlations for CG2-specific differences ran in the opposite direction (Supplemental Fig. 5). We also performed an EWCE test on this snRNA-seq data. The EWCE results showed CG2 enriched for neurons from layer 4 (FDR<0.05) and layer 5 (FDR<0.05) and CG3 enriched for neurons from layer 2 (FDR=0.006), layer 3 (p=0.04, FDR=0.40), and layer 4 (FDR=0.002, Fig. 1D). Based on these tests, we defined CG2 as representing deep-layer neurons (deepN) and CG3 as upper-layer neurons (upperN). At this juncture, the gene expression profiles for the five CGs were now garnered from the bulk brain tissue of donors with and without SCZ, i.e., upperN, deepN, a mixture of astrocytes and endothelial cells (ast_endo), oligodendrocytes (oli), and a mixture CG of astrocytes, microglia, and endothelial cells (mic_ast_endo) (Fig. 1E).

Results of this swCAM deconvolution analysis were replicated with another sample-wise deconvolution method, bMIND^30^. We applied bMIND to our tested data and identified expressions of five CGs with similar composition to those identified by swCAM. We used the Spearman correlation test to examine the similarity between CG-specific expressions estimated by swCAM and bMIND. The correlation coefficients were 0.85±0.03 (FDR<0.05) across corresponding CGs (Supplemental Fig. 6A). The showed that our deconvolution results were robust to different methods.

swCAM deconvolution results were replicated on a second RNA-seq dataset of 605 brain samples^31^. We detected five CGs in the replication dataset. (Supplemental Fig. 7). Through marker gene enrichment tests and EWCE test, we found the five CGs in these data were upperN (FDR of layer 2 neuron <0.05, p value of layer 1 neuron = 0.03), deepN (FDR of layer 4 neuron and layer 5 neuron <0.05), astrocyte (FDR<0.05), a mixture of microglia and endothelial cells (FDR<0.05) and a CG showing oligodendrocyte expression with non-significant p value (p =0.14). The dominant CGs such as upperN and deepN CGs were consistently identified.

### Generation of eQTLs for CGs

eQTL mapping was used to examine the genetic regulation of CG-specific expression. We included 72 additional brain samples from individuals with bipolar disorder (BD) to increase the sample size. Since we regressed out the affection status, the inclusion of BD patients did not interfere with the eQTL mapping results. The same five CGs were still detected by swCAM deconvolution after adding the BD samples. After removing known and hidden covariates in the CG-specific transcriptome, we focused on local eQTLs (cis-eQTLs), searching for any associations between individual expression of genes with common variants within ±1 Mb of the gene body region. In total, we identified 1,878,815 genome-wide significant eQTLs (FDR q < 0.05). After subtracting shared eQTL across all CGs, we observed more eQTL signals in neuronal CGs (both of upperN and deepN) than in glial CGs (Fig. 2A). The number of single nucleotide polymorphisms of eQTL (eSNPs) of neuronal CGs (n=302,608) was more than that of eSNPs of oligodendrocyte CGs (n=80,229). No eQTL survived multiple testing corrections in mixture CGs of astrocyte and endothelial cell data. We annotated the genomic regions for eSNPs with Ensembl annotations using ANNOVAR software^32^. The majority of eSNPs were located in intergenic and intronic regions within non-coding RNAs (Fig. 2C).

**Fig. 2.**
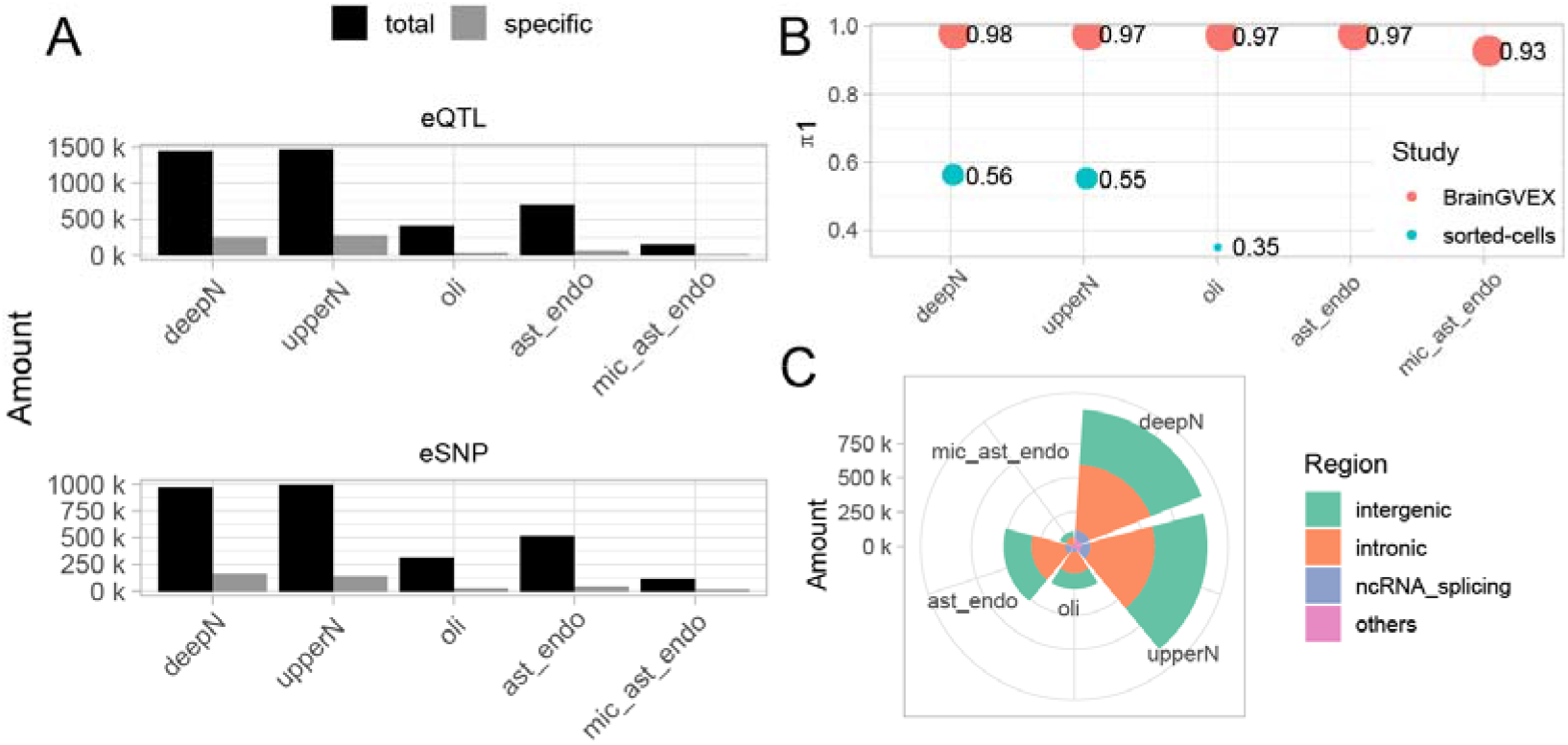
Summary of cell-group-specific eQTLs. (A) The number of eQTLs and eSNPs detected in each cell group (CG) (FDR corrected p value<0.05). (B) The similarity between CG-specific eQTLs, bulk brain tissue eQTLs, and single-cell eQTLs, evaluated by π1 values. (C) Genomic annotation of eSNPs. Ast_endo: mixture CG of astrocyte and endothelial cells, deepN: deep-layer neuron CG, upperN: upper-layer neuron CG, mic_ast_endo: mixture CG of microglia, astrocyte, and endothelial cells, oli: oligodendrocyte CG.

To examine the replication rate of eQTLs of the three CGs enriched for singular cell types (deepN, upperN, and oli), we compared our results to a sorted-cell study^33^, which profiled the gene expression and genotype of neurons (n =42) and oligodendrocytes (n =36). Using the same eQTL mapping pipeline described earlier, we calculated the eQTLs using these replication data. From this small sorted-cell dataset, we could identify 644 eQTLs for neurons and 190 eQTLs for oligodendrocytes (FDR q<0.05). We used π1 from Storey’s method^34^ to calculate the proportion of replicated eQTLs. We applied this method to the nominal p-values of replicated eQTLs in sorted cell populations, uncorrected for multiple comparisons. We found that upperN (π1=0.56) and deepN (π1=0.55) had greater reproducibility than oli (π1=0.35). Because π1 is the estimated proportion of the true alternative hypothesis, it has loose criteria for power and p-values. We used replication rate to quantify the replications, which was calculated by the ratio of replicated eQTLs among eQTLs in sorted populations with the criteria FDR<0.05. We found 29% of the neuron eQTLs were specifically replicated in neuronal CGs and 15% of the oligodendrocyte eQTLs were replicated in all CGs (Supplemental Fig. 8).

We compared our eQTL results with CG-specific eQTLs identified by bMIND. We found our results were well replicated by bMIND, with the π1 >0.98 for all CGs (Supplemental Fig. 6B). We also compared our results with eQTLs from bulk brain tissue samples from BrainGVEX. CG-specific eQTLs showed high reproducibility in the BrainGVEX dataset (π1 >0.97 in four CGs, π1= 0.92 in mic_ast_endo, Fig. 2B).

### Neuronal CGs and a mixture CG containing astrocyte and endothelial cells explain the risk heritability of SCZ

To associate brain cell type with SCZ, we used CG-specific eQTLs to assess the SCZ GWAS risk heritability. We hypothesized that if a CG is associated with SCZ, their eQTLs will explain greater GWAS heritability. To test this hypothesis, we applied stratified linkage disequilibrium (LD) score regression^35^ (sLDSC) on GWAS summary statistics and CG-specific eQTLs to partition risk heritability from summary statistics. The SNP-based heritability analysis showed neuronal CGs and a CG mixture containing astrocyte and endothelial cells significantly enriched for risk heritability of SCZ (Fig. 3A). The upper-layer neuron CG explained 19.24% of heritability with 14% SNPs (adjusted p value=2.97e-04, Fig. 3B). The deep-layer neuron CG explained 19.03% of heritability with 14% SNPs (adjusted p value=4.81e-04). Mixture CG of astrocyte and endothelial cells explained 12% of heritability with 8% of SNPs (adjusted p value=1.04e-03). These results suggest the regulatory regions in the neuronal CGs and a mixture CG of astrocyte and endothelial cells have a higher proportion of SCZ common variants than that in oligodendrocyte CG and a CG mixture containing microglia, astrocyte, and endothelial cells. We also compared enrichment score in bulk tissue eQTLs and CG-specific eQTLs. We found the enrichment scores at bulk level (1.29±0.07) and cell levels (1.42±0.17) were very close (Fig. 3A).

**Fig. 3.**
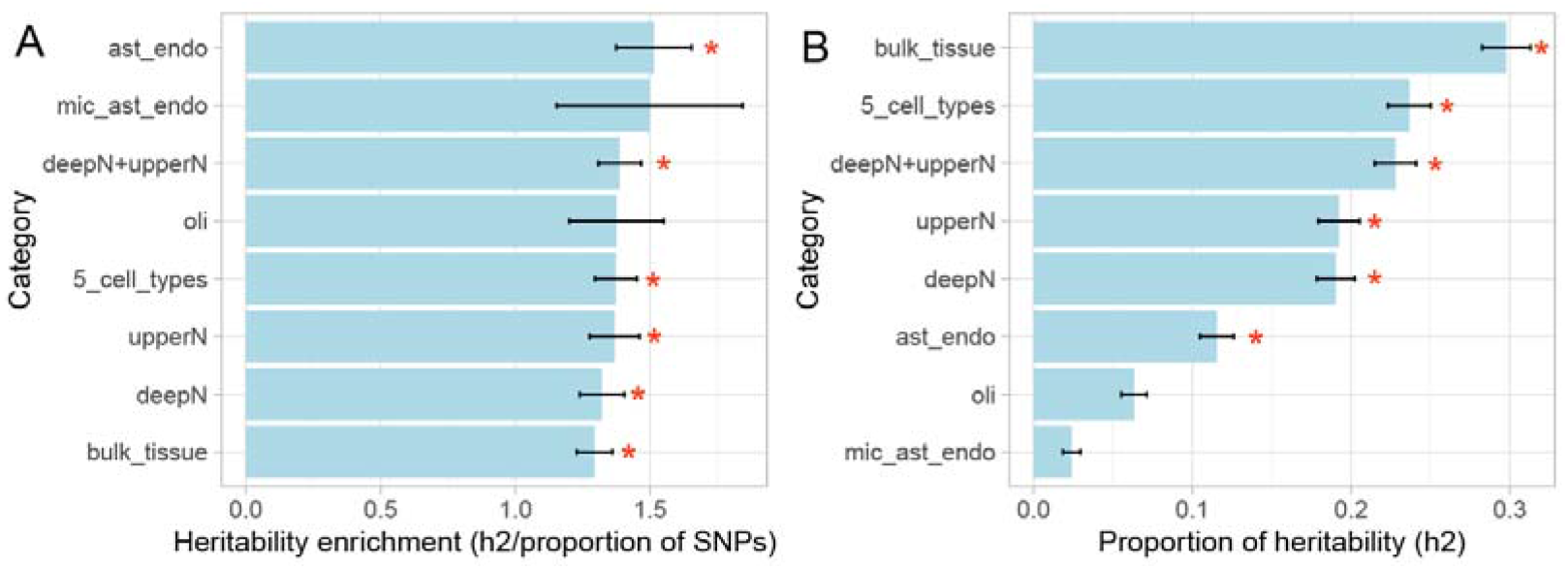
Genetic risk heritability enrichment in cell groups. GWAS heritability was partitioned in five estimated cell groups (CGs). The SNPs of CG-specific eQTLs (FDR corrected p value<0.05) were cataloged in annotated files in sLDSC. (A) the SCZ risk heritability enrichment in different categories (calculated by the proportion of heritability (h2) / proportion of SNPs) and (B) the h2 explained by different categories. The asterisk denotes a corrected p value <0.05 in the enrichment test. ast_endo: mixture CG of astrocyte and endothelial cell, deepN: deep-layer neuron CG, upperN: upper-layer neuron CG, mic_ast_endo: mixture CG of microglia, astrocyte, and endothelial cells, oli: oligodendrocyte CG.

### Decreased proportion of upper-layer neuron CGs from donors with SCZ

To determine if the proportion of each CG was altered in the brains of patients with SCZ, we estimated the proportion of CGs for each sample using CAM. The proportion of upperN was reduced by 2.76% in patients with SCZcompared with controls, and the mixture CG of mic_ast_endo increased by 0.8% in patients with SCZ (Wilcoxon signed-rank test, p value<0.05, Fig. 4A). Other CGs did not show significant change. The proportional change in upper-layer neurons was statistically significant in a permutation test (p<0.001, Supplemental Fig. 9).

Based on our testing for genetic enrichment, we hypothesized that the changes in CG proportion could arise from genetic risk variants. We used two strategies to test this hypothesis. First, we performed QTL mapping using CG proportions as the phenotype, but we did not detect any significant CG proportion-QTL signals. Second, we used polygenic risk score (PRS) as a measure of genetic risk burden for each individual and tested the correlation between PRS and CG proportions. We observed weak negative correlations between SCZ PRS and the upperN proportion and the p value did not survive multiple testing (r=-0.1, p value=0.04, FDR=0.1, Fig. 4B).

**Fig. 4.**
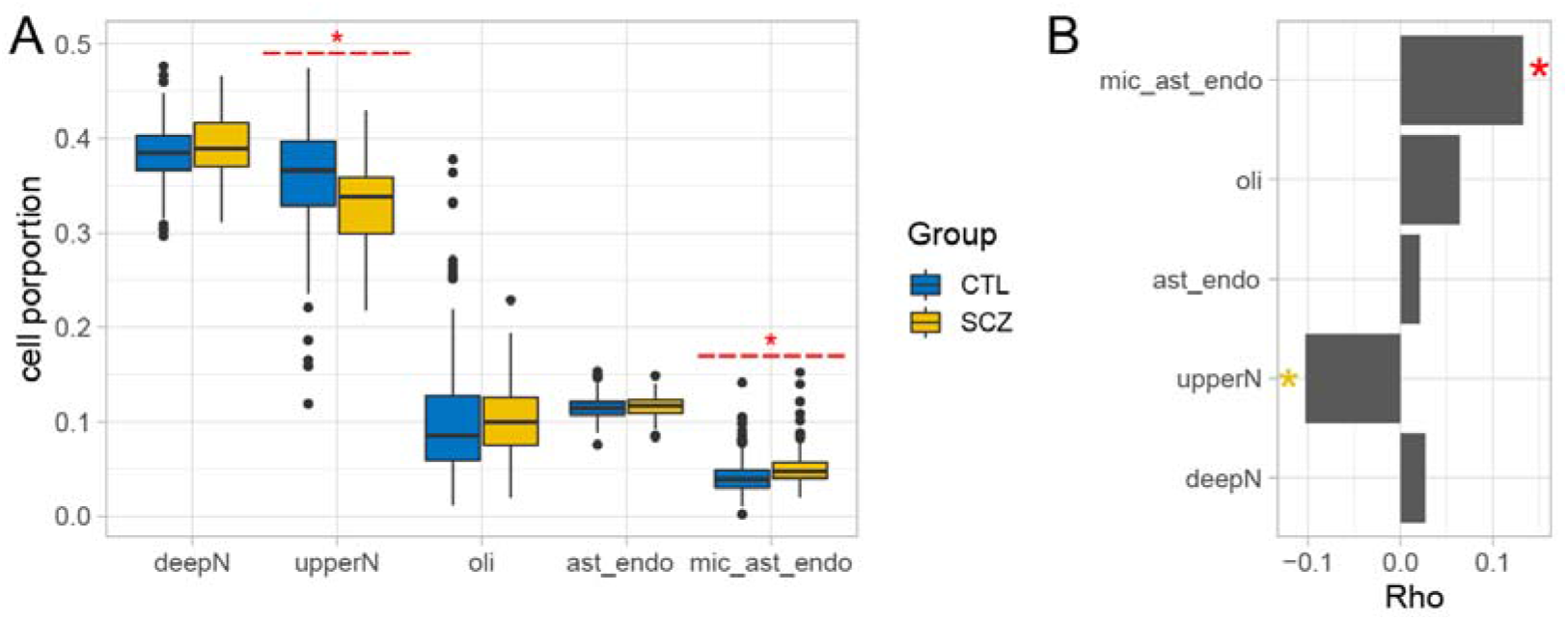
Proportion per cell group and association with SCZ. (A) Proportions of the five cell groups (CGs) for donors with and without SCZ, estimated with the CAM method. The asterisk denotes corrected p-values of the Wilcoxon rank-sum test <0.05. (B) Spearman correlation between polygenic risk score (PRS) and CG proportion. The red asterisk denotes corrected p-values of correlation test <0.05 and the yellow asterisk denotes nominal p value <0.05. Ast_endo: mixture CG of astrocyte and endothelial cells, deepN: deep-layer neuron CG, upperN: upper-layer neuron CG, mic_ast_endo: mixture CG of microglia, astrocyte, and endothelial cells, oli: oligodendrocyte CG.

### CG-specific differential gene expression in SCZ

We further analyzed CG-specific differential expression comparing individuals with and without SCZ using Wilcoxon signed-rank test with significance by FDR q <0.05, (Supplemental Table 2). We compared our results with differential expression results from bMIND and observed significant consistency (Fisher’s exact test p value<0.05, Supplemental Fig. 6C). We found substantial overlap in differentially expressed genes (DEG) across five CGs, with 330 upregulated genes and 344 downregulated genes shared across five CGs (Supplemental Fig. 10). The number of shared DEGs across all CGs was greater than the number of CG-specific DEGs, except for upregulated genes in CG mic_ast_endo, with 367 genes. Approximately 33% of the genes (n=14,865, Gencode v19) were affected in at least one CG in the SCZ cohort. The two neuronal CGs possessed greater transcriptional changes than the three glial CGs, (fold change: 0.20 in neuronal CGs versus 0.09 in glial related CGs, p value<2.2e-16, Kolmogorov–Smirnov test). We performed permutation tests to determine the empirical probability of transcriptional changes in CGs. We found that the fold changes observed in all CGs were significant (p<0.001). Larger fold changes in neuronal CGs than that in the glia CG were also significant in permutations (p<0.001, Supplemental Fig. 11).

We used Gene Ontology (GO) enrichment to annotate the function of DEGs (Table 1), focusing on the functions of dysregulated genes in neuronal groups. A total of 1,537 downregulated DEGs were detected in the upperN. These were enriched for protein folding (FDR q= 3.20e-09) and antigen processing (FDR q= 8.01e-05). The 1,407 downregulated DEGs within deepN were enriched for positive regulation of cell proliferation (FDR q= 2.24e-06) and trans-synaptic signaling (FDR q=4.10e-06). Upregulated genes in both the upperN (n=229) and deepN (n=223) GCs were enriched for carboxylic acid breakdown (FDR q_deepN_=1.04e-06, FDR q_upperN_ =6.93e-08).

**Table 1.**
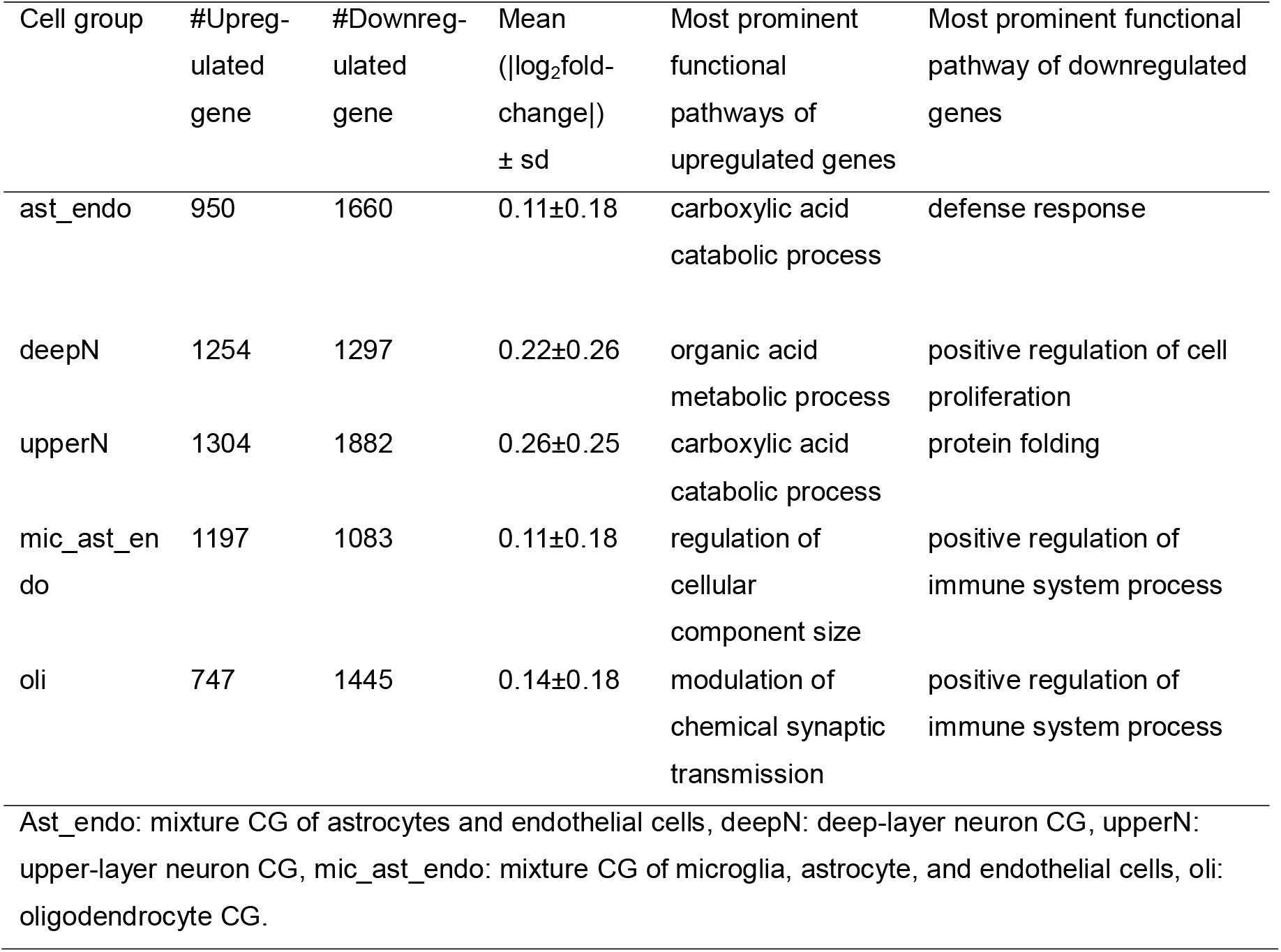
Summary of cell group-specific differential gene expression.

### Transcriptome-wide association analysis identified SCZ-associated genes in all CGs

To determine which genes had CG-specific dysregulated expression driven by genetics, we performed a TWAS analysis using the CG-specific eQTLs. We identified 51 SCZ risk genes across 43 genetic loci (adjusted p value<0.05, Supplemental Table 3). We compared our TWAS results to the 193 SCZ risk genes identified in bulk tissue by TWAS in Gandal et al^16^. Twelve genes in our results had been previously identified in Gandal’s results. Our analysis attributed these 12 genes to all 5 CGs: deepN (*CYP17A1-AS1, AS3MT, CSPG4P11, EMB*), upperN (*CYP17A1-AS1, AS3MT, CSPG4P11, EMB, GATAD2A, NAGA, RP11-73M18.8, TYW5*), ast_endo (*RP11-73M18.8, CYP17A1-AS1, EMB*), oli (*SNX19, AC011330.5, CSPG4P11, EMB*), and mic_ast_endo (*BTN3A2, TOM1L2, CSPG4P11*). Additionally, 39 CG-specific TWAS genes are novel to bulk tissue-based TWAS.

Out of the 51 TWAS genes, 17 genes are in deepN and 30 in upperN. Eight genes existed in both subtypes (*CYP17A1-AS1, AS3MT, PIGB, GOLGA6L9, CSPG4P11, CNPPD1, EMB, ENDOG*). Twenty-two genes were uniquely identified in upperN (*DIABLO, C12orf65, RP11-73M18.8, STRC, AP3B2, WHAMM, SLX1B, ARL17B, GATAD2A, RPS5, ADAM15, RP1-130H16.16, WBP2NL, NAGA, ALMS1, TYW5, ITIH4, TMEM110, SFMBT1, NDUFAF2, BTN2A1, DNPH1*). Nine genes were identified in the deepN only (*CWF19L2, MORN3, TIE1, RBBP5, CBR3, RFTN2, RP11-10L12.4, REEP2, STK4*). Twenty-four TWAS genes are specific to glial CGs (9 from ast_endo, 10 from oli and 5 from mic_ast_endo).

### Co-expression network analysis refined CG-specific biological processes associated with SCZ

To investigate the coordinated expression changes in each CG, we performed weighted gene correlation network analysis^36^ (WGCNA) on CG-specific expressions (Supplemental Table 4). For this study, we identified *CG-specific modules*, a group of genes showing concordant variation across samples in one specific CG. Modules of upperN and deepN were highly consistent. In total, we identified eight CG-specific modules out of the 149 detected modules. Seven of the eight showed either differential expression of eigengenes in SCZ or enrichment of SCZ GWAS signals. These seven modules were defined as “SCZ-associated modules” (Table 2).

**Table 2.** Cell group-specific co-expression modules.

| Cell group | Module name | Trait association | GWAS enrichment | Representative GO term (FDR<0.05) | Top 3 hub genes |
| --- | --- | --- | --- | --- | --- |
| ast_nedo | Ast_endo_ME28 | - | SCZ2014, Education, IBD | RNA splicing | SEC62, PPIG, PNN |
| neuron | neuron_ME28 | SCZ↑ | SCZ2014, Education | neuronal stem cell population maintenance | TSC22D1, ENTPD3, SYNE1 |
|  | neuron_ME29 | - | SCZ2014, Education | SWI/SNF superfamily-type complex | WDR48, STOML2, OS9 |
|  | neuron_ME30 | SCZ↑ | SCZ2014, Education, Neuroticism | carbohydrate transmembrane transport | FKBP5, SORT1, WIPF3 |
|  | neuron_ME32 | - | SCZ2014, Education | negative regulation of actin filament polymerization | GRIA4, CUX2, EPHX4 |
|  | neuron_ME27 | - | SCZ2014, Education, Neuroticism, BD2016 | regulation of dendritic spine morphogenesis | RIMS3, NGEF, LZTS3 |
| mic_ast_endo | mic_ast_endo_ME25 | - | - | Defense response to virus | RSAD2, DDX58, PSME2 |
| oli | oli_ME14 | SCZ↓ | - | Ensheatment of neuron | RNASE1, GPR37, PLP1 |

Ast\_endo: mixture CG of astrocyte and endothelial cells, deepN: deep-layer neuron CG, upperN: upper-layer neuron CG, mic\_ast\_endo: mixture CG of microglia, astrocyte, and endothelial cells, oli: oligodendrocyte CG.

The neuronal CGs shared five of the seven SCZ-associated modules (Fig. 5). The eigengenes of two modules showed upregulated expression in SCZ: neuron_ME28 (log2FC=0.02, FDR=6.8e-03) and neuron_ME30 (log2FC=0.03, FDR=6.3e-07). These two were enriched in the pathways of neural stem cell maintenance and carbohydrate transmembrane transport (FDR q<0.05). The other three modules showed only enrichment of GWAS signals (neuron_ME32, neuron_ME27, and neuron_ME29)and were enriched within pathways of regulation of actin filament polymerization, regulation of dendritic spine morphogenesis, and SWI/SNF superfamily-type complex, respectively (FDR q<0.05).

**Fig. 5.**
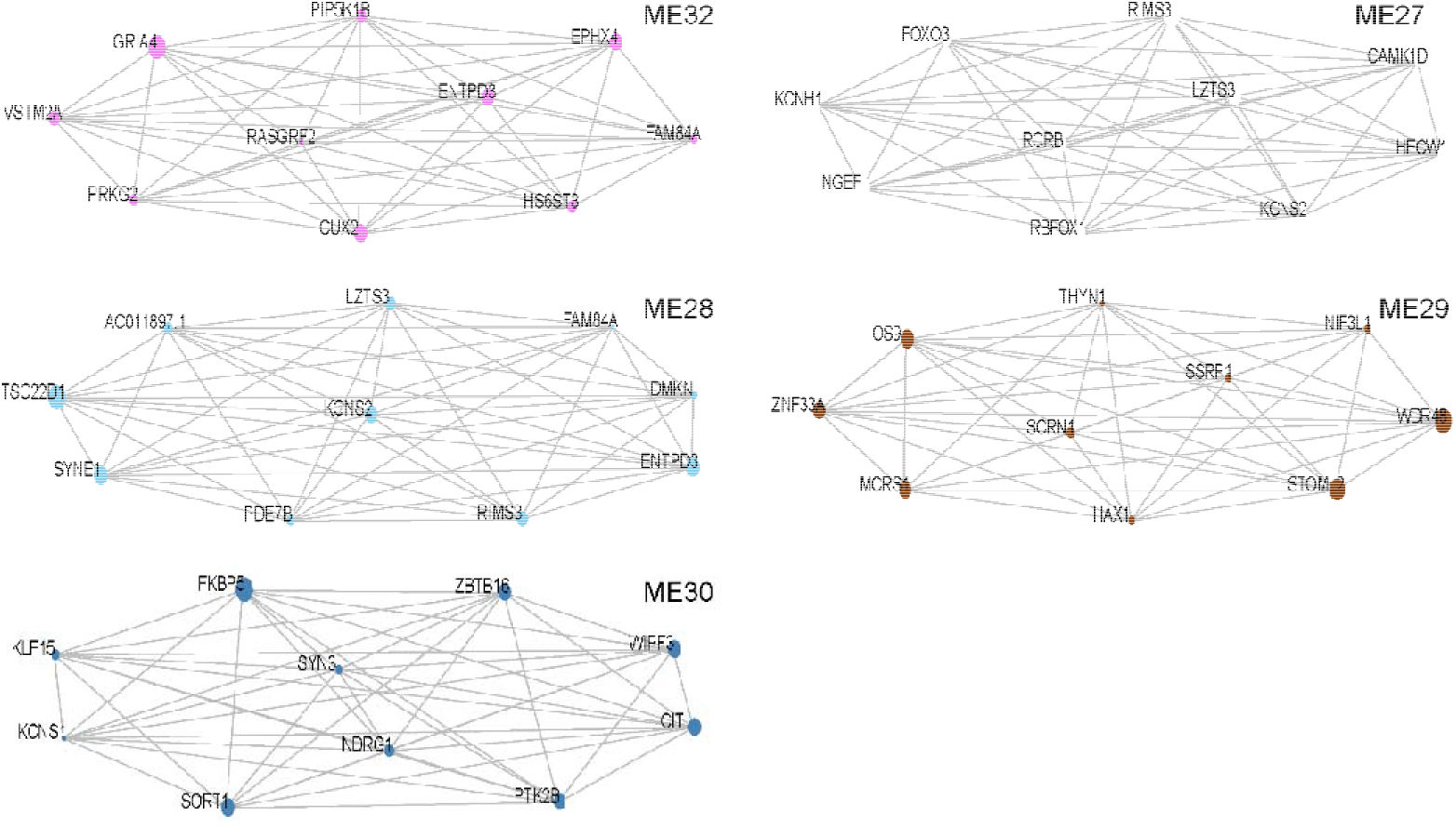
Neuron-specific co-expression modules related to SCZ. Top 10 genes ordered by module membership (kME, i.e., correlation between gene expression and module eigengene) are shown for each neuron-group-specific module.

The other two SCZ-associated modules were specific to glial CGs. In the mixed CG of astrocytes and endothelial cells, we identified ast _endo_ME28, enriched for SCZ GWAS signals, and involved with the RNA splicing pathway (FDR q<0.05). In the oligodendrocyte CG-related module (oli_ME14), the eigengene was downregulated in SCZ (log2FC=-0.01, FDR=0.04). Member genes in this oli_ME14 were involved in neuron ensheathment (FDR q<0.05).

## Discussion

Combining CG-specific eQTL analysis and GWAS can reveal *how* (the regulation of gene expression) and *where* (within which CGs) variants contribute to disease risk. Recent studies have employed this approach identifying cell-type-specific eQTLs in blood that have heightened risk heritability for immune-related diseases^37,38^ However, cell-type-specific eQTLs for human brains are not available and are difficult to obtain. To bridge genetic findings of SCZ to brain cell types, we generated CG, a proxy of cell type, using an unsupervised deconvolution method. We then mapped CG-specific eQTLs using computationally partitioned gene expression specific to each CG per individual. Based on GWAS data, we found enrichment for risk heritability in the eQTLs of upper-layer DLPFC neuronal CGs (upperN). Furthermore, we found that the upperN CGs also displayed dramatic gene expression changes and altered cell proportions in the individuals with SCZ.

Our results are consistent with previous bulk tissue studies of human and cell-type-specific studies of mice in implicating neurons in the risk for SCZ. In a case-control study of bulk brain tissues, Gandal et al. showed the neuron-specific co-expression module to be downregulated in SCZ^16^. Skene et al. calculated the neuronal specificity of the human-homologous gene in mouse scRNA-seq data and correlated this specificity to risk heritability enrichment within the genetic region^39^. They found that four neuron subtypes in the mouse brain (i.e., medium spiny neurons, pyramidal cells in hippocampal CA1, pyramidal cells in the somatosensory cortex, and cortical interneurons) are associated with SCZ. Our study expands upon those findings, confirming that a subtype of neurons, the group of upper-layer neurons in human DLPFC, are associated with SCZ.

Our study provides genetic support for an association between the upper-layer neurons of the DLPFC and SCZ. This is supported by the enrichment of GWAS signals, dysregulated gene expression, and decreased proportions within the upperN CGs. Previous research has shown upper-layer neurons of the DLPFC are over-represented in humans compared with other species,^40^ and to be actively engaged in the delayed activity of working memory^41^, the impairment of which is a major feature of SCZ. Some phenotypes of upper-layer neurons within the DLPFC have been reported to be associated with SCZ in terms of cell shape^42^, dendritic abnormalities^43,44^, and candidate mRNA expression^45^. However, no genetic evidence that we are aware of to date has pinpointed upper-layer neurons as having a role in SCZ risk. In this study, we observed that GWAS risk variants contribute to SCZ by altering gene expression within the upper-layer neurons. Our coexpression network analysis and TWAS analysis further narrowed down specific genes and pathways. Therefore, we suspect that previously reported phenotypic changes^42–46^ within the upper-layer neurons of patients with SCZ may be rooted in these genetic variants and their related gene expression regulation.

Our findings of SCZ-associated changes in upper-layer neurons were validated by multiple lines of experiments from other studies. In a recent snRNA-seq study of cortical layers from seven SCZ patients and 11 controls, the selective vulnerability of supragranular layer neurons was revealed in SCZ in terms of dramatic transcriptomic changes and cell proportion changes^47^. A decreased proportion of upper-layer neuron subtypes was also observed in mouse model SCZ-predisposing 22q11.2 microdeletion^48^ and human postmortem brains^49,50^. Moreover, the functional mechanism of a differential expressed gene in upper-layer neuron CYFIP1 (log2FC=−0.13, FDR=0.0004) has been discovered in the mouse cortex. It has been reported that CYFIP1 deficiency causes upper-layer neurons to present in deep layers and deep-layer neurons to present in upper layers^51^. Moreover, the changes in upper-layer neurons also have been noted in association with another psychiatric disorder, autism, based on snRNA-seq study^18^. In summary, this study strengthened the findings of upper-layer neuron dysfunction in SCZ and provided more detailed cell-subtype evidence based on a relatively large brain collection. It provides strong support to previous studies of animal models, cell models, and brain transcriptome studies with limited samples.

Our findings suggest the SCZ-associated changes in upper-layer neurons may occur in neuron development. Neuron development in human brain includes proliferation, differentiation, neuron migration and synaptic formation. This process is coupled with substantial transcriptome changes^52^. We found that the DEGs in upperN were enriched in the pathways of cell proliferation and trans-synaptic signaling, which has previously been suggested as involved in SCZ risk. For example, the disrupted proliferation of neurons has been reported in primate and murine models of SCZ^53^. Five TWAS genes in upperN are related to neuronal development as well: *AS3MT, CSPG4P11, ADAM15, STRC, and WHAMM.* For example, *AS3MT* participates in neural stem cell differentiation and is a risk gene for SCZ^54^. *CSPG4P11* is also an eQTL gene in fetal brains, and its expression has also been linked to SCZ risk^55,56^. *ADAM15, STRC*, and *WHAMM* function in cell-matrix adhesion, which is linked to neurodevelopment and SCZ^57,58^. Therefore, our findings support the hypothesis that the disturbed gene expression in neurodevelopment is involved in SCZ molecular pathology. While our study cannot provide causality of these genes in SCZ pathology, future functional experiments in cells and useful animal models could potentially confirm the functional mechanism of genes.

While swCAM yielded reproducible unsupervised deconvolution results, particularly for neuronal CGs, it also has its limitations. The CGs estimated by swCAM represent groupings of cells with similar transcriptional profiles. Correspondingly, swCAM has relatively poor resolution in separating rare cell types. Glial subtypes with small compositions, such as microglia, have been particularly challenging. As single-cell data are accumulating, a hybrid model integrating a suitable reference may boost swCAM’s resolution. Despite these constraints, swCAM enhances the merit of existing bulk-tissue data by deriving expression for major CGs. With this method, it is possible to determine whether disease-related gene expression is a feature of specific CGs or a pattern of composite CGs. Furthermore, gene networks and genetic regulatory relationships can emerge that may help to ferret out hidden clues to the molecular mechanisms of complex psychiatric disorders.

## Conclusion

Of five clusters of cell types estimated by gene expression *in silico*, the upper-layer neurons within the DLPFC display distinctive enrichment of risk heritability and transcriptional changes in the brain tissue of donors with SCZ. Furthermore, decreased proportion of upper-layer neurons relative to al cell groups was observed within the SCZ brain. The uniquely dysregulated genes and pathways associated with upper-layer neurons, particularly those related to neurodevelopment, should be targeted in future experimental studies and quantitative analysis of SCZ to further understand the molecular mechanism of SCZ.

## Materials and methods

### Brain tissues

The PsychENCODE/BrainGVEX project has 427 postmortem DLPFC samples, including healthy controls, BD and SCZ patients. Brain samples originated from the Stanley Medical Research Institute (SMRI) and the Banner Sun Health Research Institute (BSHRI). Detailed information can be found at https://www.synapse.org/#!Synapse:syn4590909.

### RNA sequencing

Total RNA was isolated from brain tissue at the University of Illinois at Chicago and at the University of Chicago with the Qiagen miRNeasy mini kit. Quality of RNA samples was measured with an Agilent Bioanalyzer RNA 6000 Nano assay kit and all samples had a RIN score of 5.5 or greater. All total RNAs were processed into stranded, rRNA-depleted libraries for sequencing in an Illumina HiSeq2000. Libraries were triplexed per lane to reach 40 million paired-end reads per library. Details of library preparation and RNA sequencing can be found in our previous study^16^.

### Data processing

Data processing included three steps: reads processing, quantification, and post-quantification processing. Fastq files underwent adapter removal using cutadapt (https://pypi.python.org/pypi/cutadapt), after which the resulting adapter-trimmed fastq files were checked for quality using FastQC (http://www.bioinformatics.babraham.ac.uk/projects/fastqc/). A subset of 10,000 reads estimated the insert mean size and standard deviation for use with Tophat. STAR was used to align trimmed reads to the GENCODEv19 reference (modified to include artificial ERCC RNA ExFold spike-in sequences). BAM files were sorted in samtools (v1.3). Expression level was then calculated using RSEM (v1.2.29). Quality control metrics were calculated using RNA-SeQC (v1.1.8), featureCounts (v1.5.1), PicardTools (v1.128), and Samtools (v1.3.1). The quantified fragments per kilobase of transcript per million mapped reads (FPKM) were used for downstream analysis. Mitochondrial genes, pseudoautosomal genes, and genes with FPKM fewer than 1 in more than 75% of samples were dropped. We calculated the distance between samples and removed samples with a z-score of normalized connectivity to other samples lower than −2. After filtering, 341 samples and 14865 genes were retained for subsequent analyses. Linear regression removed the effect of covariates including age, sex, RIN, PMI, brain bank, batches, and principal components of sequencing statistics (seqPC). The seqPCs was composed of the top 29 principal components analyzed from the sequencing statistics. Covariates were selected by multivariate adaptive regression splines (MARs).

### Genotype processing

Genotypes were called from three platforms, including 277 samples by whole genome sequencing, 257 samples by Psych v1.1 beadchips, and 137 samples by Affymetrix 5.0 450K. Genotypes were imputed by Michigan Imputation Server using HRC (r1.1 2016) EUR samples as references from each platform. Variants with R squared less than 0.3 or HWE (Hardy-Weinberg Equilibrium) p-value less than 1e-6 were removed. We used DRAMS software^59^ to correct mixed-up data IDs for the three platforms based on relationships across various omics data. We had 251 samples genotyped by whole genome sequencing and beadchip. The genotypes called from the two platforms were highly consistent (r= 0.986). Genotypes of the three platforms were combined, with discordant genotypes marked as missing. Polygenic risk scores were calculated using PRSice (2.1.4) with GWAS summary data from Ripke et al^2^ as base data set.

### Sample-specific deconvolution

To isolate CGs *in-silico*, we developed a novel sample-wise deconvolution technique (swCAM^26^), applying it to the processed RNA-seq data. The original CAM framed the deconvolution of mixture expression to solve the blind source separation problem (sBSS), which was accomplished by nuclear norm regularization. The basic assumption in the deconvolution was

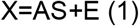

which is the mixture sample being the weighted sum of the CG-specific expression S. In the classic deconvolution method, S is a common matrix for all mixture samples. However, each sample may have its sample-specific S_i_:

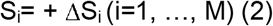

The associated sample-specific BSS model is given by

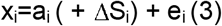

If some samples have a group pattern in a particular source or some molecules’ expression are more highly correlated in one particular source, the associated samples may share a similar pattern of ΔS_i_, leading to the following swCAM objective function:

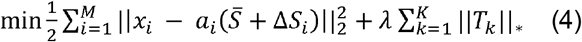

Where T_k_ consists of the kth column in all ΔS_i_, representing between-sample variability in source K; ||T_k_||* is the nuclear norm of T_k_. λ is the regularization parameter of nuclear norms. We solved equation (4) using quadratic programming and the alternating direction method of multipliers (ADMM). Code of swCAM can be accessed at https://github.com/Lululuella/swCAM.

We used CAM to estimate the CG proportion in an unsupervised manner. The processed expression data was the input of CAM. The minimum description length (MDL) algorithm was used in CAM^21^ to automatically determine the number of CGs (K). MDL is a widely-adopted and consistent information theoretic criterion in model selection. The shortest MDL denotes the best model. Shortest MDL was shown when K=5 for the data. The proportions of five CGs and processed RNA-seq data were entered into swCAM to infer the CG-specific expression for each sample. To obtain the top ctDEGs and thereby differentiate the CGs, Wilcoxon signed-rank test determined the difference of expression in each CG (log2FC>2, FDR<0.05).

### Processing of single-cell datasets

Three single-cell datasets provided further confirmation of CG identities^28,29,60^. We downloaded raw counts from frontal cortices and used the Seurat package (v2.0) for data processing^61^. We filtered out the following: genes expressing in less than half of the smallest CGs, cells with unique feature counts (i.e., >5,000 and <200), and cells with mitochondrial counts >5%. The data were then normalized by library size and log-transformed. The cell identities from the original studies were used for cell clustering. The averaged expression was calculated to represent gene expressions for the specific cell type.

### Processing the ROSMAP dataset for replication

The synapse website (10.7303/syn3388564) provided a replication dataset with 640 postmortem human DLPFCs from the Religious Orders Study and Memory and Aging Project (ROSMAP)^31^. Gene expression was measured by RNA-seq. The downloaded raw expression matrix was quantified in FPKM. After quality control, the data retained 19,144 genes and 605 samples. Expression was log2-transformed and normalized by quantile normalization. The confounder effects from age, sex, diagnosis, sequencing batch, study, ancestry, education year, and PMI were removed by linear regression.

### Cell group identification

We used two strategies to annotate the CGs; gene sets enrichment and correlation with scRNA-seq data. In the identification of five general CGs, the marker genes of five major cell types were collected from Zhang et al^27^ and the marker genes for inhibitory neurons and excitatory neurons were collected from Lake et al^63^. Fisher’s exact test was used to test the enrichment of marker genes in CGs. Meanwhile, Spearman correlation testing with scRNA-seq datasets^28,60^ was used to measure similarities between CG expression and that of known cell types. To identify genes expressed in gradient cortical layers, we downloaded RNA-seq data across human DLPFC layers from the He et al. study^64^. Gene expression was set as a linear regression function for DLPFC layers (from layer 1 to layer 6). Genes with coefficient >0 and FDR <0.05 were named “deep-layer genes.” Those with coefficient <0 and FDR <0.05 were named “upper-layer genes”. scRNA-seq data used for correlation testing with neuronal CGs was downloaded from https://portal.brain-map.org/atlases-and-data/rnaseq, which provided neuronal expression for the multiple layers of human middle temporal gyri^29^. Spearman correlation test was performed on layered neuronal expression and that of CG2 and CG3.

Expression Weighted Cell Type Enrichment^65^ (EWCE) test was used as a replication of CG annotations. The purpose of EWCE is to determine the chance of a target gene list having higher expression in a specific cell type than a random gene list. The ctDEGs identified in each CG were used as target lists. Two snRNA-seq data sets from frontal cortex^17,28^ and one snRNA-seq data set from six cortical layers^29^ were used as tested data. The test was repeated 10000 times. The p values were corrected by the Benjamini-Hochberg method.

### CG-specific differential expression

Differential expression analysis was conducted in each CG with the Wilcoxon rank-sum test. The p values were corrected by FDR. Genes with FDR q value <0.05 were identified as differentially expressed genes. To calculate the empirical probability of observed fold changes, we conducted a permutation test as follows. In each CG, we permutated the case/control labels and calculated the fold changes of case-control differential expression. The permutation test was repeated 1000 times.

### Pathway analysis

Gene ontology enrichment tests for biological processes, molecular function, and cellular components were performed using gProfileR^66^, with *p*-values FDR-corrected.

### CG-specific eQTL mapping

To identify CG-specific eQTLs in the brain, we performed cis-eQTL mapping using package FastQTL^67^. eQTL mapping was performed independently for each CG. We used CG-specific expression estimations from swCAM as phenotype data. The phenotype data were gene expression matrixes for five CGs (five matrixes formatted in genes * samples). A cis-window was defined for genes in each CG as 1 mega base up- and down-stream of the gene body. Each CG expression matrix was adjusted for hidden covariates using PEER^68^ with FDR-corrected *p*-values. eQTLs with FDR <0.05 were retained.

To compare CG-specific eQTLs with sorted-cell eQTLs and bulk brain eQTLs, we used Storey’s Qvalue package^34^ and replication rate. The proportion of true associations (π1) was estimated for significant CG-specific eQTLs in the sorted-cell eQTLs/BrainGVEX DLPFC eQTLs. With the distribution of corresponding *p* values for the overlapping eQTLs, we calculated π0, i.e., the proportion of true null associations based on distribution. Then, π1 = 1 - π0 estimated the lowest bound for true-positive associations. The replication rate is the ratio of replicated eQTLs with FDR<0.05 among all significant eQTLs in sorted cell populations.

### Heritability estimation with GWAS summary statistics

Partitioned heritability was measured through LD Score Regression v1.0.0^69^, identifying enrichment of GWAS summary statistics among CG-specific eQTLs by accounting for LD. CG-specific eQTL annotation files were created by eSNPs detected in each CG. The annotation file was produced by marking all HapMap3 SNPs that fell within the eQTL annotations. LD-scores were calculated for SNPs within the file (LD window of 1cM). The LD reference panel was downloaded from 1000 Genomes European Phase Three. SNP LD-scores were interwoven in the computation of partitioned heritability (proportion of h^2^). Enrichment for each annotation file was calculated by the proportion of heritability explained by each annotation file divided by the proportion of SNPs falling in that annotation file. Enrichment *p* values were then corrected by multiple testing.

### Estimation and case-control comparison of cell proportions

To compare the CG composition in SCZ patients and healthy controls, we used CAM to estimate the proportion of each CG. Wilcoxon rank-sum test was used to compare the CG proportions between SCZ patients and healthy controls. FDR correction was performed on the p values. To determine the empirical probability of proportion changes in upper-layer neuron CG, we permutated the case/control labels and calculated the case-control proportion changes in upper-layer neuron CG. This process was repeated 1000 times.

### Transcriptome-wide association study

We performed TWAS analysis on the GWAS summary statistics^70^ and the CG-specific eQTLs using the TWAS/FUSION package^9^. To identify risk genes with evidence of genetic control, we used genome-wide complex trait analysis to estimate cis-SNP heritability h2g (±1MB window around gene body). We identified genes with significant h2g (nominal p-value <0.05), which were used to calculate the SNP-based predictive weights per gene. Using the FUSION package, we calculated five-fold cross-validation of five models of expression prediction and evaluated the prediction models for accuracy. The five models are best cis-eQTL, best linear unbiased predictor, Bayesian sparse linear mixed model [BSLMM], and elastic-net regression, LASSO regression. The model with the largest cross-validation R^2^ was chosen for downstream association analyses. TWAS statistics were calculated using different cell-group weights, LD SNP correlations from the 1000 Genomes European Phase 3 reference panel. TWAS association *p* values were FDR-corrected and were corrected for multiple CGs.

### Gene co-expression analysis

We constructed the gene co-expression network for each CG using Weighted Gene Co-Expression Network Analysis (WGCNA) on deconvoluted CG-specific expression data. A correlation matrix was calculated for the genes, which was then weighted after a scale-free topology was approximated. The following weighted powers were chosen for the respective CGs: 12 for ast_endo and deepN, 14 for upperN, mic_ast_endo, and oli. The signed adjacency, biweight midcorrelation, and blockwiseModules parameters were selected to build the network. Other parameter settings included the following: mergeCutHeight= 0.1, minModuleSize= 30, pamStage=FALSE, and deepSplit=4.

### Associating the gene modules with SCZ

Two strategies were used to test the association between gene modules and SCZ. First, we constructed a linear model in which the dependent variable is the module eigengene across all samples and the independent variable is the disease state for the samples (FDR-adjusted *p* value<0.05). Second, we tested the correlation between GWAS significance and the module membership (kME), which is the expression correlation between each module member gene and the module eigengene. GWAS significance of each module member gene was calculated by MAGMA v1.06^71^ using the European subset of the 1000 Genomes as a reference panel for LD. An annotation step was performed first in which GWAS significant SNPs were mapped to genes, based on the presence of a GWAS significant SNP in the region between a gene’s start and stop sites +/− 10kb. A competitive gene-set analysis was then performed using sets defined by gene co-expression modules. Resulting *p* values were FDR-corrected for multiple comparisons. Modules meeting any of the two criteria above were defined as “SCZ-associated modules”.

### Gene module preservation test

To test the similarity among the CG-specific modules, we tested preservation among the modules detected per CG using Z_summary_. The network-based preservation test generated two types of statistics: 1) density-based preservation statistics to determine whether the nodes in the reference network were still highly connected in the test network; 2) connectivity-based preservation statistics to determine whether the connectivity pattern in the reference network was preserved in test network. We applied Z_summary_ test by Peter Langfelder et al^72^, in the formula

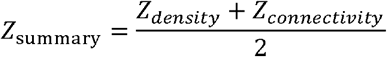

Z_density_ summarizes density preservation statistics, and Z_connectivity_ summarizes connectivity-based statistics.

## Data availability

Access to the raw BrainGVEX data is controlled by the NIMH Repository and Genomics Resources (NRGR), https://www.nimhgenetics.org/. Instructions can be found in the PsychENCODE Knowledge Portal: https://www.synapse.org/#!Synapse:syn4921369. Deconvoluted data can be reached at http://lbpg.upstate.edu/module_search/. Source data are provided with this paper.

## Code availability

Scripts used in the data analysis for this manuscript can be found at GitHub: https://github.com/RujiaDai/CellSpecificAnalysis. Code for swCAM can be found at https://github.com/Lululuella/swCAM.

## Acknowledgments

This work was supported by NIH grants 1U01 MH103340-01, 1R01ES024988, and New York State Empire Innovation Program (to C.L.) and National Natural Science Foundation of China (NSFC) grants 31970572 and 31871276, National Key Plan for Scientific Research and Development of China grant 2016YFC1306000, and Innovation-Driven Project of Central South University grants 2020CX003 (to C.C.). We thank the PsychENCODE consortium and other data contributors (De Jager et al, Habib et al, Zhang et al, Darmanis et al, Lake et al, Mendizabal et al, Hodge et al, He et al) for their data support. We thank James Liu for contributions in web site design and data visualization. We gratefully acknowledge the families of the brain donors, without whom this work would not have been possible.

## Author contributions

Data processing: R.D., S.L., Y.J. Bioinformatics analysis: R.D., S.L., Y.C, J.D. Algorithm development: L.C., C.W., G.Y., Y.W. Web design: R.D., Q.W. The conception of study design: R.D., S.L, C.C., C.L. Writing of the manuscript: R.D. Substantive revisions of the manuscript: R.D, C.L., C.C., Y.W., R.K.

## Competing interests

The authors have no competing financial interests to declare.

## Additional information

This work was honored with the Hugh Gurling Memorial Awards by the International Society of Psychiatric Genetics at XXVI World Congress of Psychiatric Genetics.

## Supplemental materials

**Supplemental Fig. 1.**
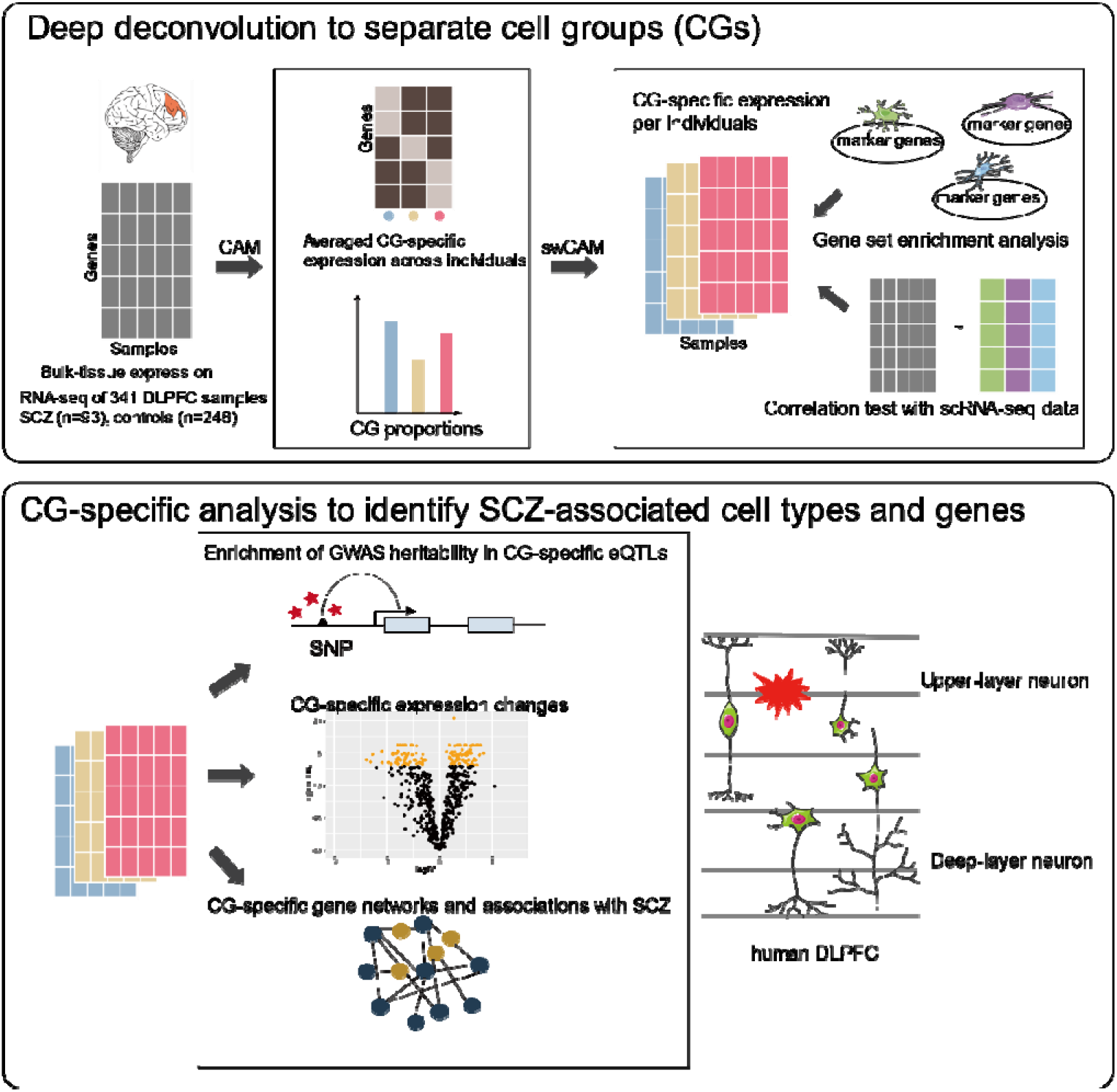
Overview of the study. Using the sample-wise convex analysis of mixtures (swCAM) deconvolution method, we extracted cell-group-specific (CG-specific) expression from RNA-seq data from bulk brain tissue of 341 samples (SCZ=93, CTL=248). To annotate the identified CGs, maker gene enrichment and correlation testing with scRNA-seq data were used. To discover SCZ-associated cell types and genes, we performed CG-specific eQTL mapping, GWAS heritability enrichment, CG-specific differential expression, CG-specific co-expression, and transcriptome-wide association study. These analyses offer a framework for discerning cell subtypes and underlying genes and pathways associated with SCZ. We found upper-layer neuron CGs in human DLPFC to be associated with SCZ.

**Supplemental Fig. 2.**
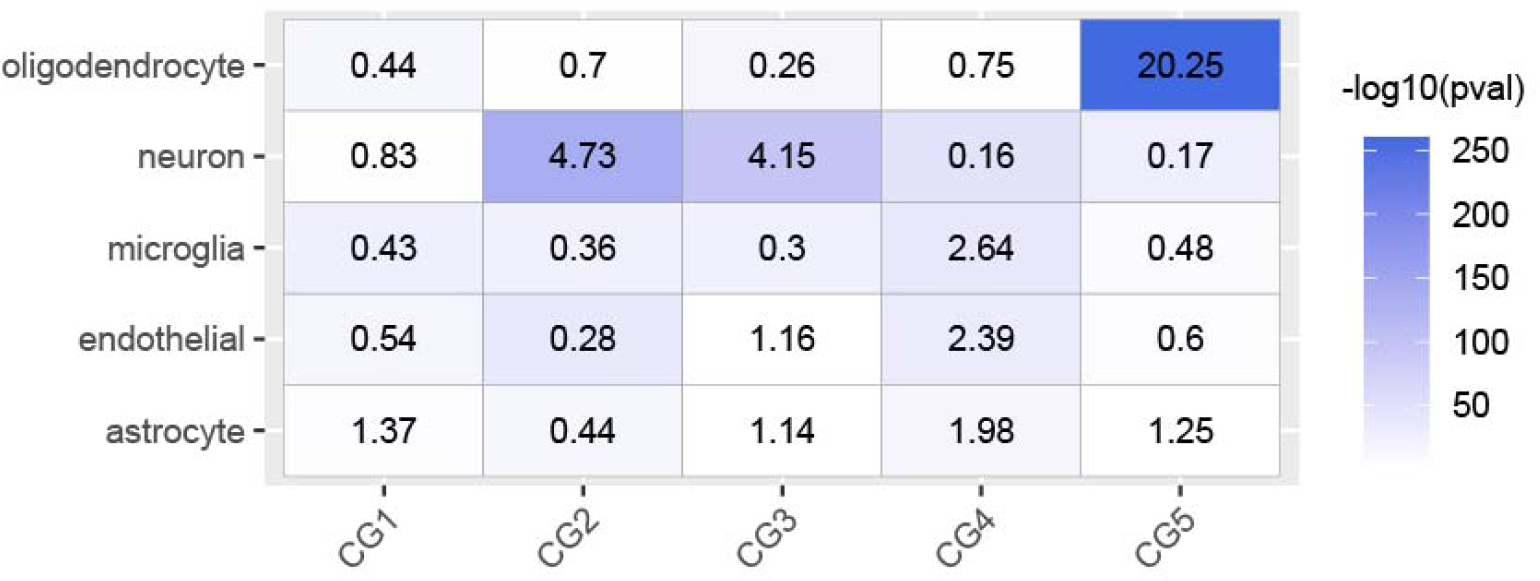
Cell-group enrichment of brain cell marker genes. Enrichment of the top cell-type differentially expressed genes (ctDEGs) in estimated cell groups (CGs) (log_2_FC>2, FDR <0.05) for human brain cell marker genes. Color denotes log_10_-scaled p values of enrichment for significant associations and text in cell denotes odds ratio (Fisher’s exact test).

**Supplemental Fig. 3.**
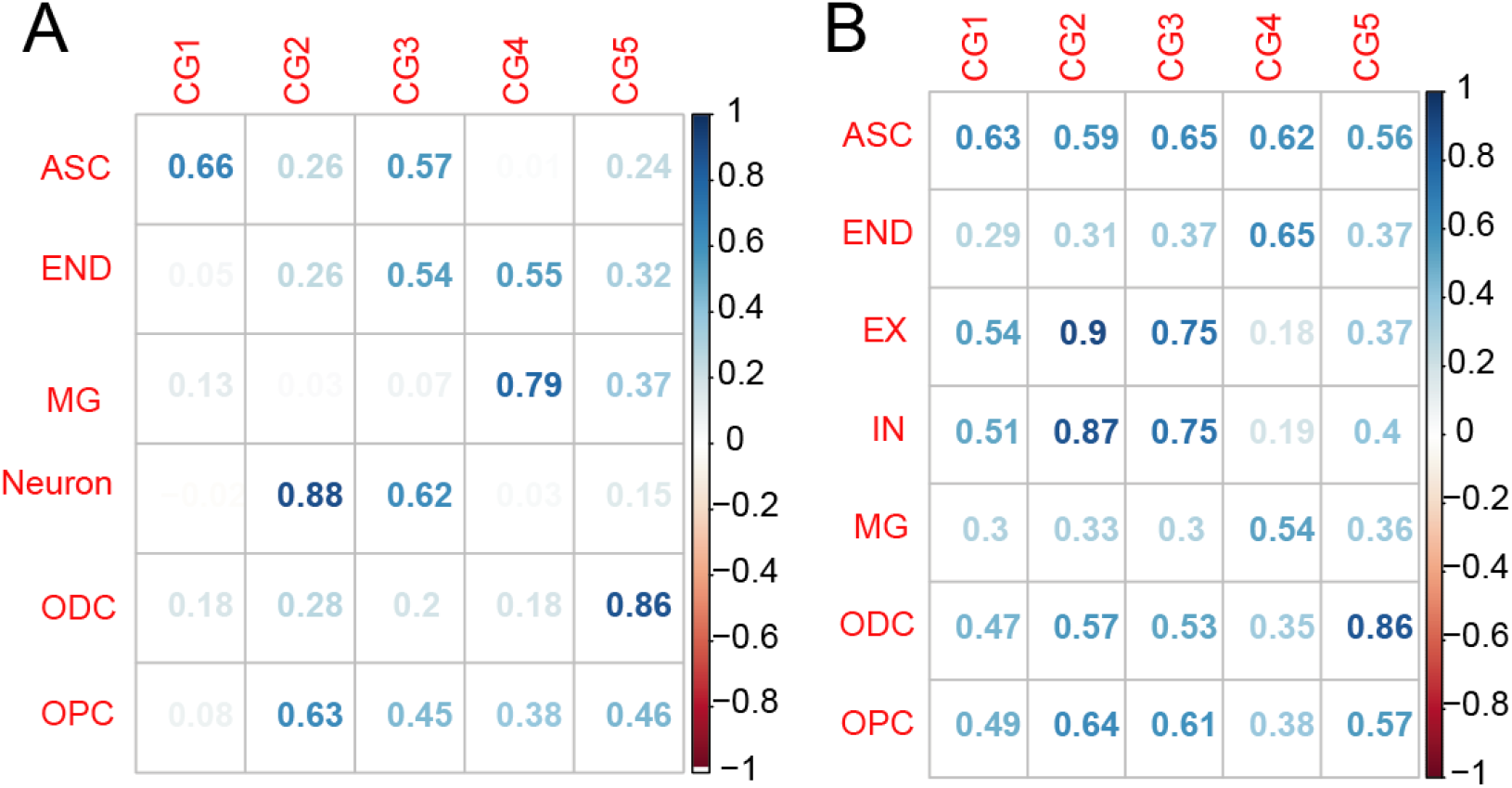
Correlations between estimated cell-group-specific (CG-specific) expression and (A) single-cell expression (Darmanis et al. 2015) and (B) single-nuclei expression (Habib et al. 2017). The Spearman correlation test was used. The number and the color both denote the correlation coefficient. ASC: astrocytes, END: endothelial cells, MG: microglia, ODC: oligodendrocytes, OPC: oligodendrocyte precursor cells, EX: excitatory neurons, IN: inhibitory neurons.

**Supplemental Fig. 4.**
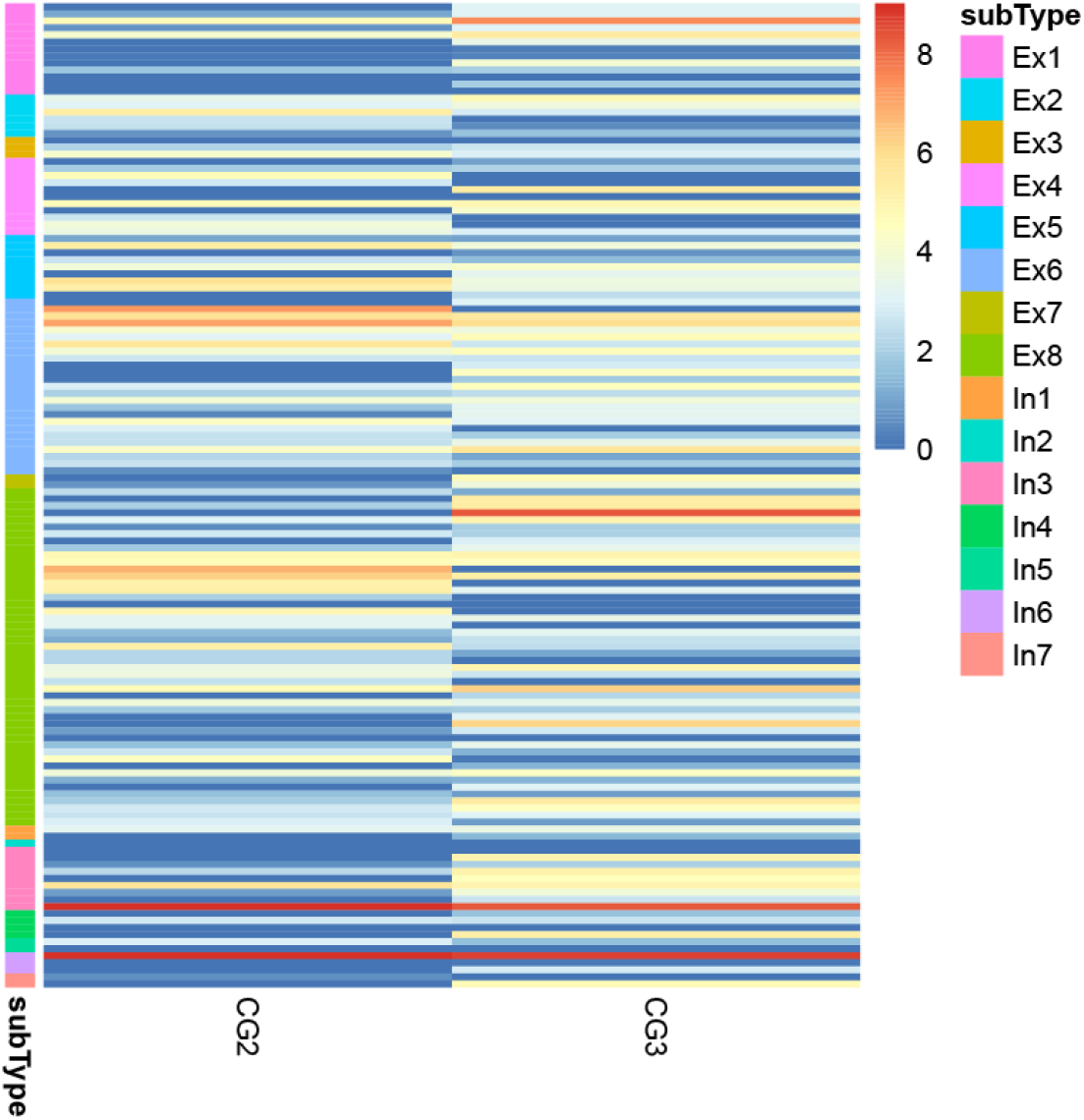
Expression of marker genes of the inhibitory and excitatory neuronal subtypes in deconvoluted neuronal cell groups (CGs). Ex: excitatory neurons; In: inhibitory neurons. The data of subtypes was from a single-nucleus study of human brains (Lake et al, 2017).

**Supplemental Fig. 5.**
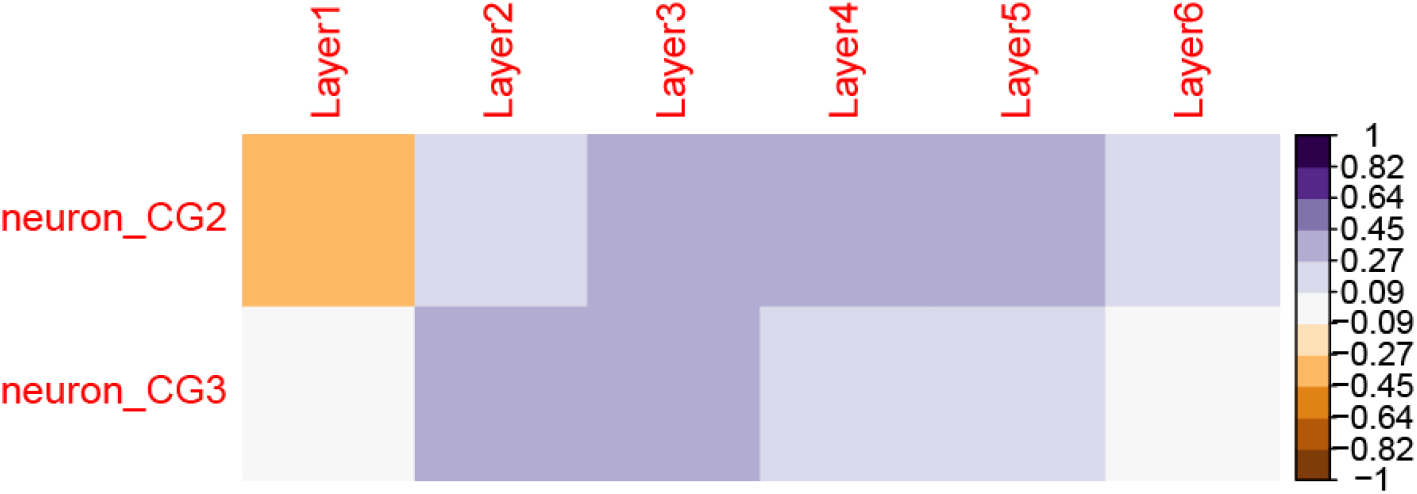
Spearman correlation between expressions of estimated neuronal cell groups (CGs) and expressions of neurons from different layers of the middle temporal gyrus.

**Supplemental Fig. 6.**
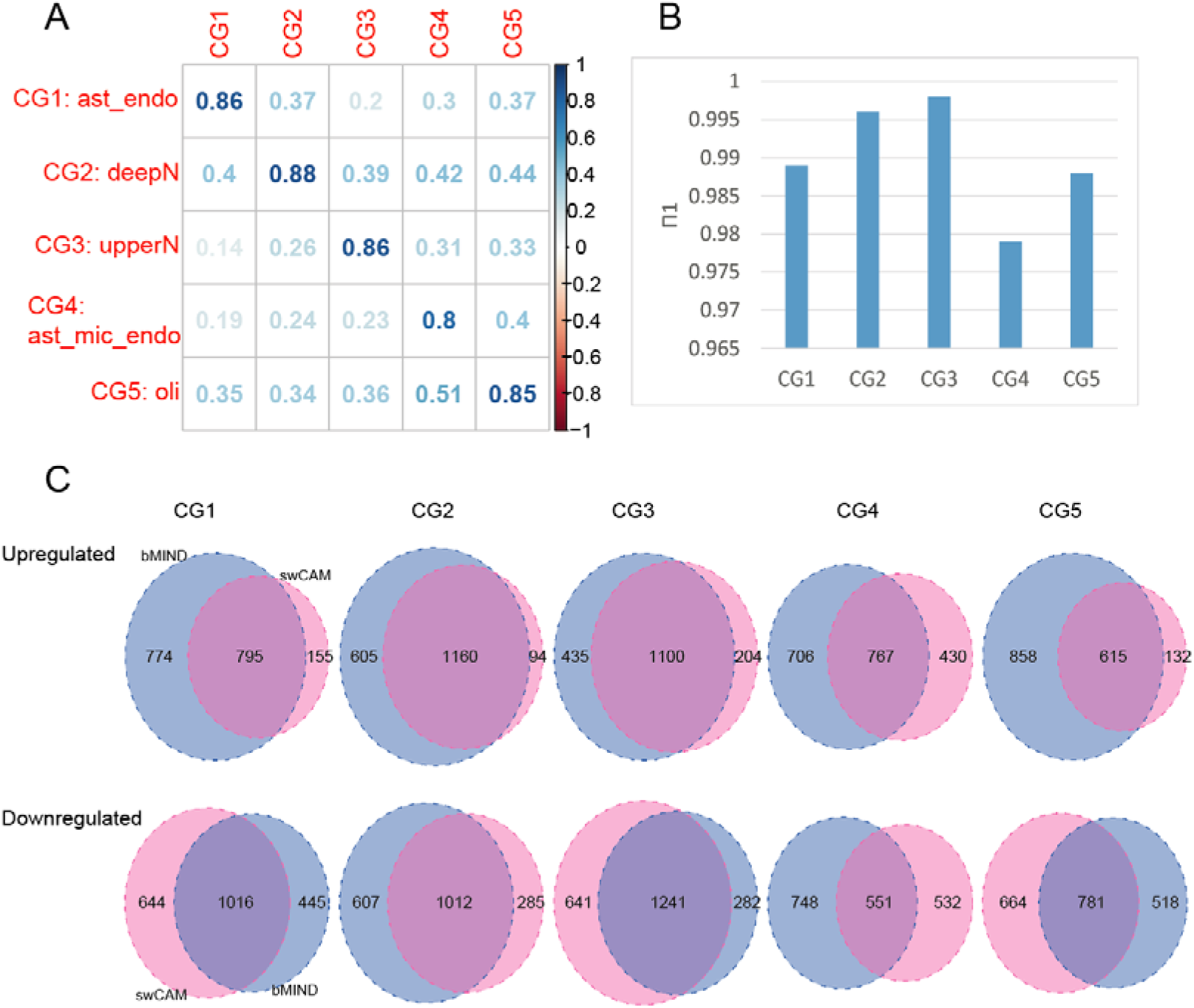
Comparison with an alternative deconvolution method bMIND. (A) Spearman correlations between CG-specific expressions estimated from swCAM (rows) and bMIND (columns). (B) Replication of CG-specific eQTLs by bMIND. (C) Comparison of SCZ differentially expressed genes identified by swCAM and bMIND.

**Supplemental Fig. 7.**
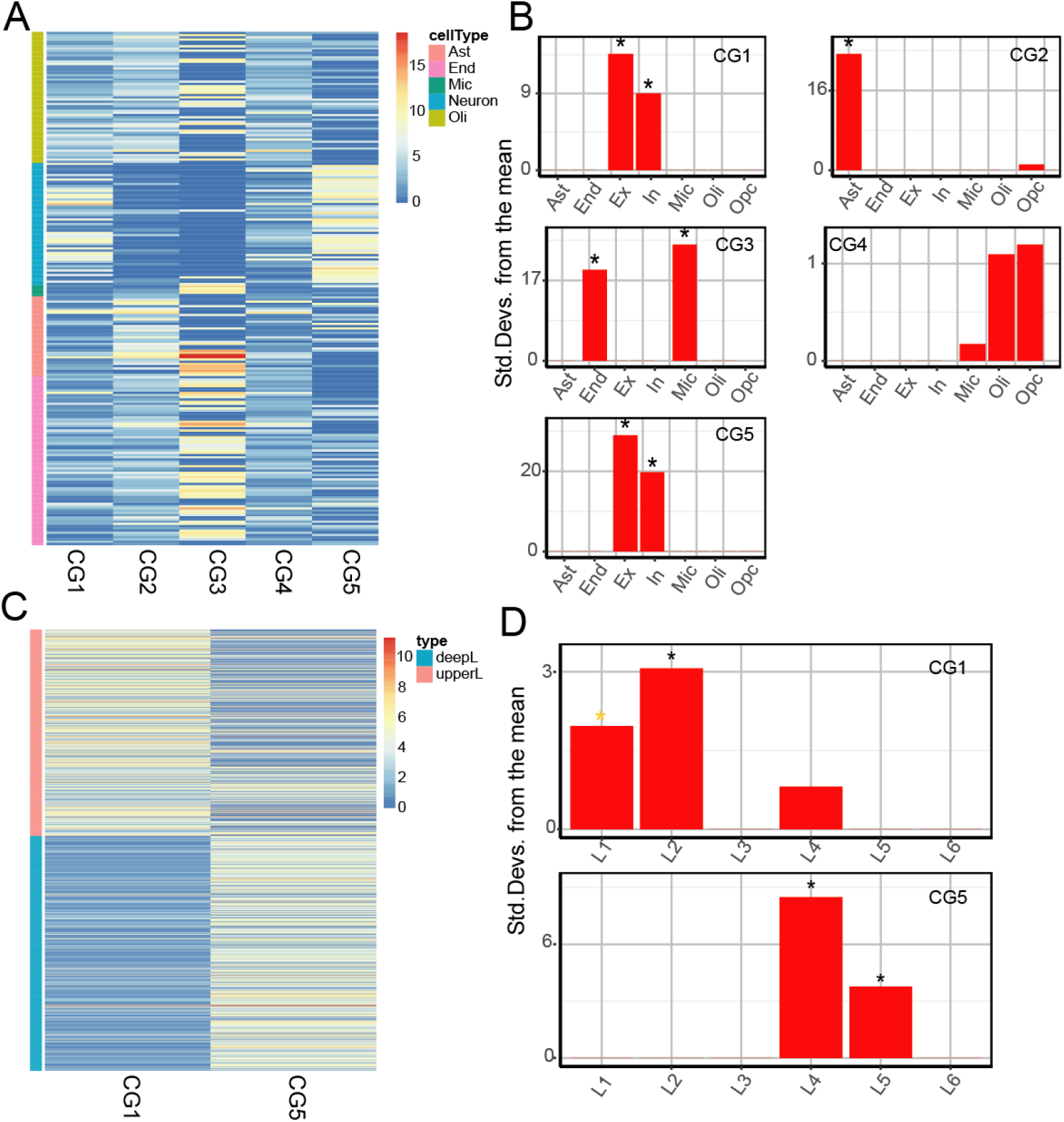
Deconvolution of replication dataset from ROSMAP project. (A) Marker gene expression in estimated cell groups (CGs). (B) To annotate the identity of CGs, an EWCE test with snRNA-seq data from human cortex was conducted. (C) Expressions of upper-layer and deep-layer genes in estimated CGs. (D) To annotate the identity of neuronal CG1 and CG5, EWCE test with snRNA-seq data of neurons from cortical layers was conducted. Ast: astrocytes, End: endothelial cells, Mic: microglia, Oli: oligodendrocytes, Opc: oligodendrocyte precursor cells, Ex: excitatory neurons, In: inhibitory neurons.

**Supplemental Fig. 8.**
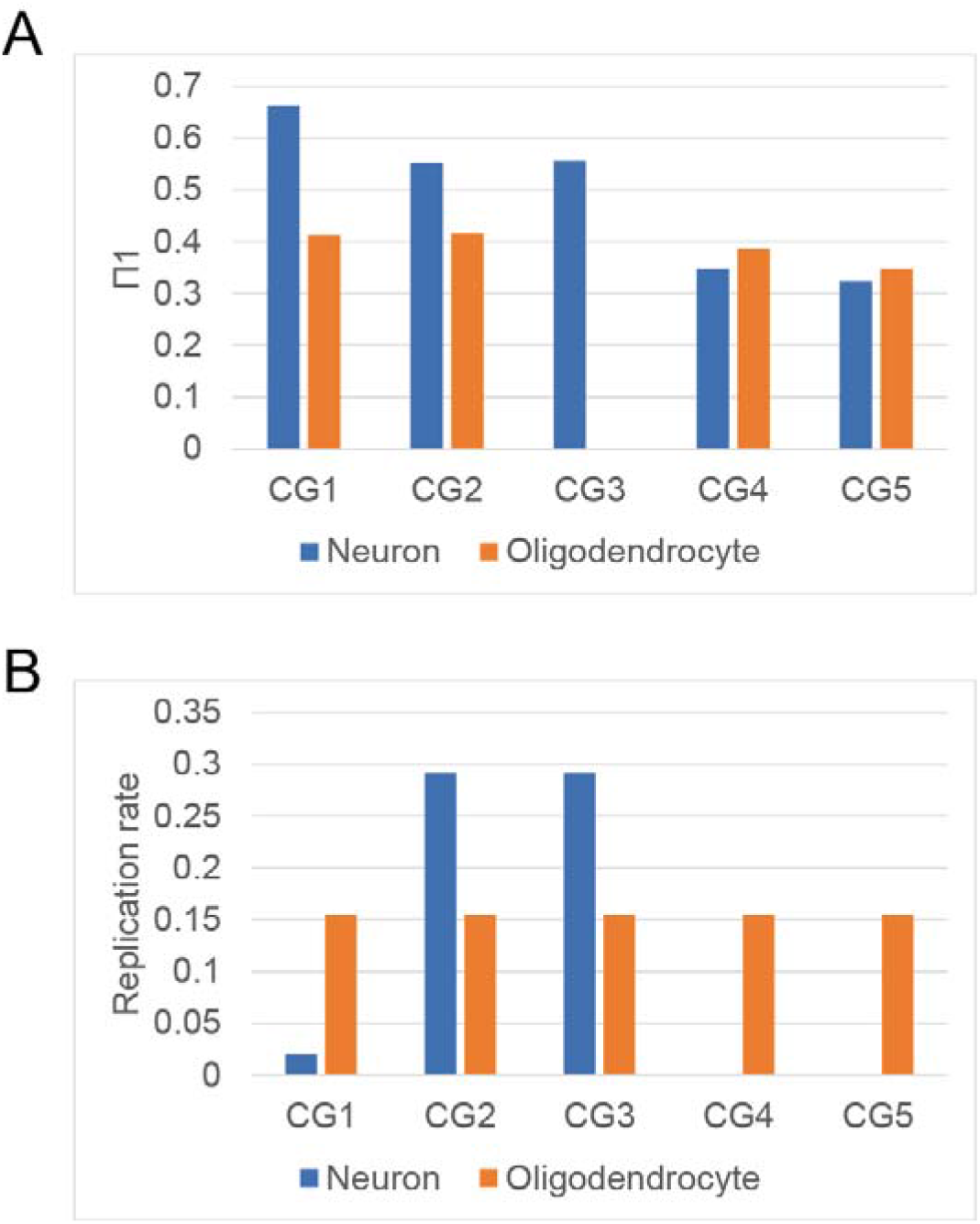
Replication of CG-specific eQTLs in eQTLs from sorted cell populations. (A) The degree of replication is indexed by Π1 and (B) replication rate. Π1 is the proportion of true alternative hypothesis of replicated eQTLs in sorted population. Replication rate is the ratio of replicated eQTLs over eQTLs in sorted populations (FDR<0.05).

**Supplemental Fig. 9.**
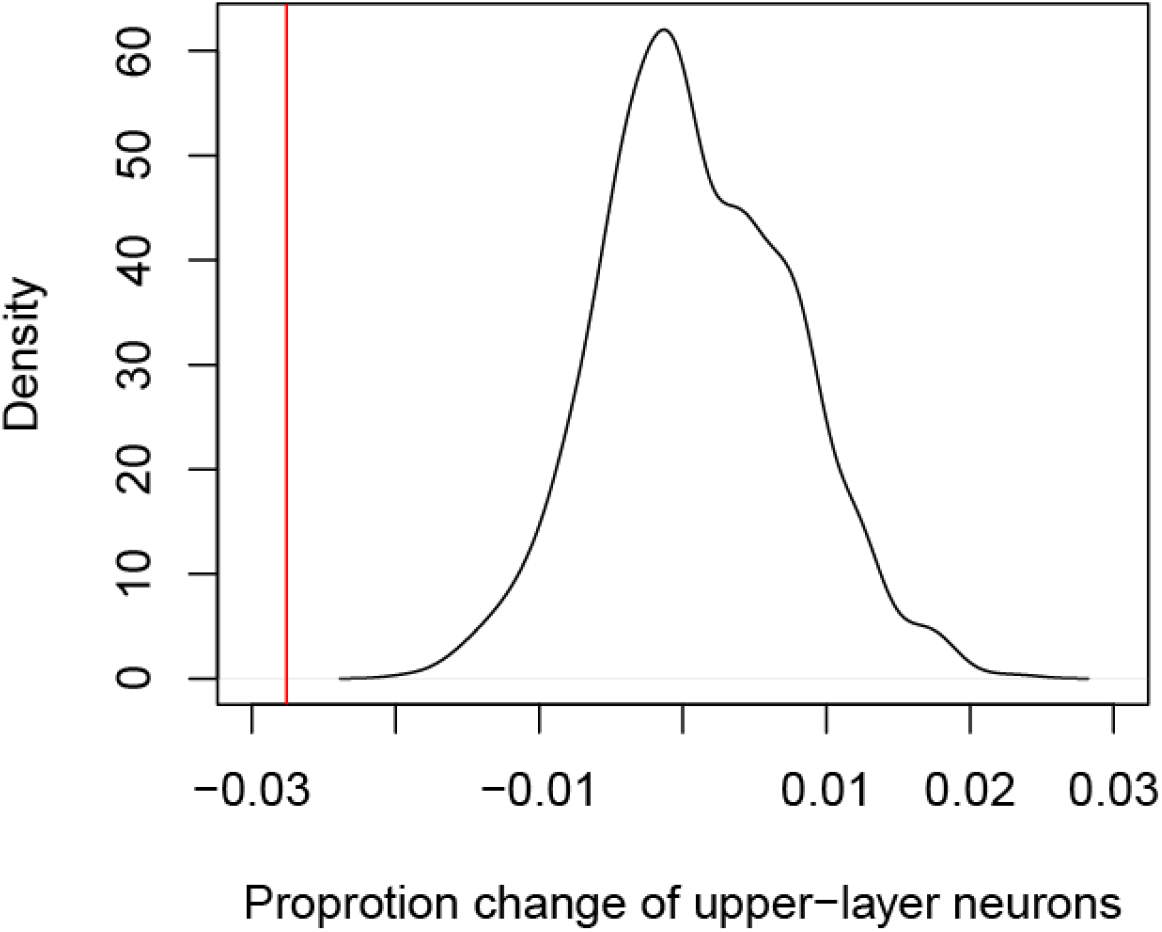
Permutation test of proportion changes observed in upper-layer neurons. The sample labels were shuffled and the proportion changes were calculated from two randomly selected groups. This process was permutated 1000 times. The black line is the distribution of proportion changes in permutations and the red line is the observed proportion changes.

**Supplemental Fig. 10.**
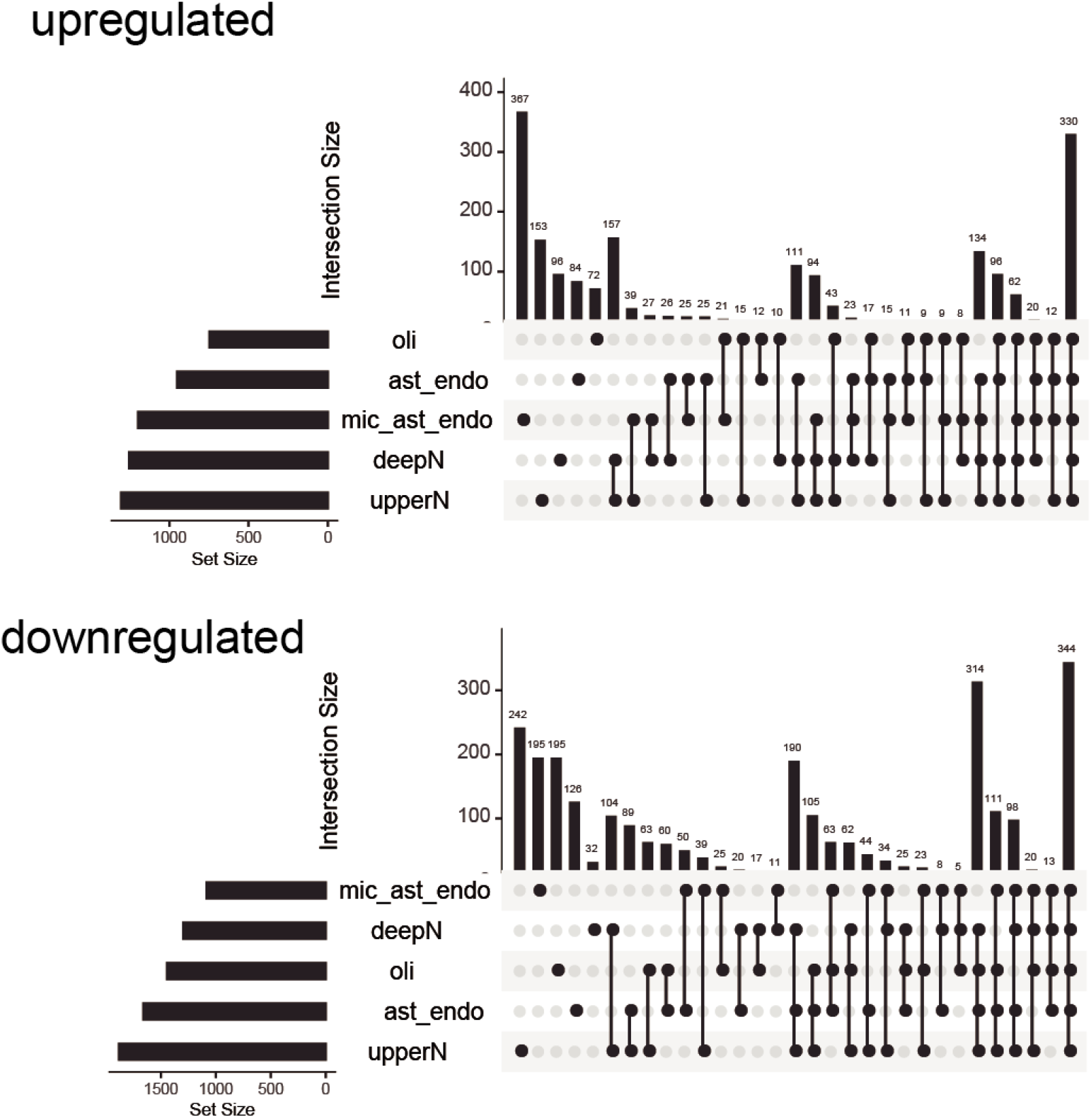
Overlap of differentially expressed genes detected in each cell group. The differential expression of genes in patients with SCZ was calculated with Wilcoxon signed-rank test (p value <0.05) per cell group (CG). The left-side bars denote the number of differentially expressed genes in each CG. The upper bars denote the intersection size between sets of differentially expressed genes. Dark connected dots on the bottom panel denote which substrates are considered for each intersection.

**Supplemental Fig. 11.**
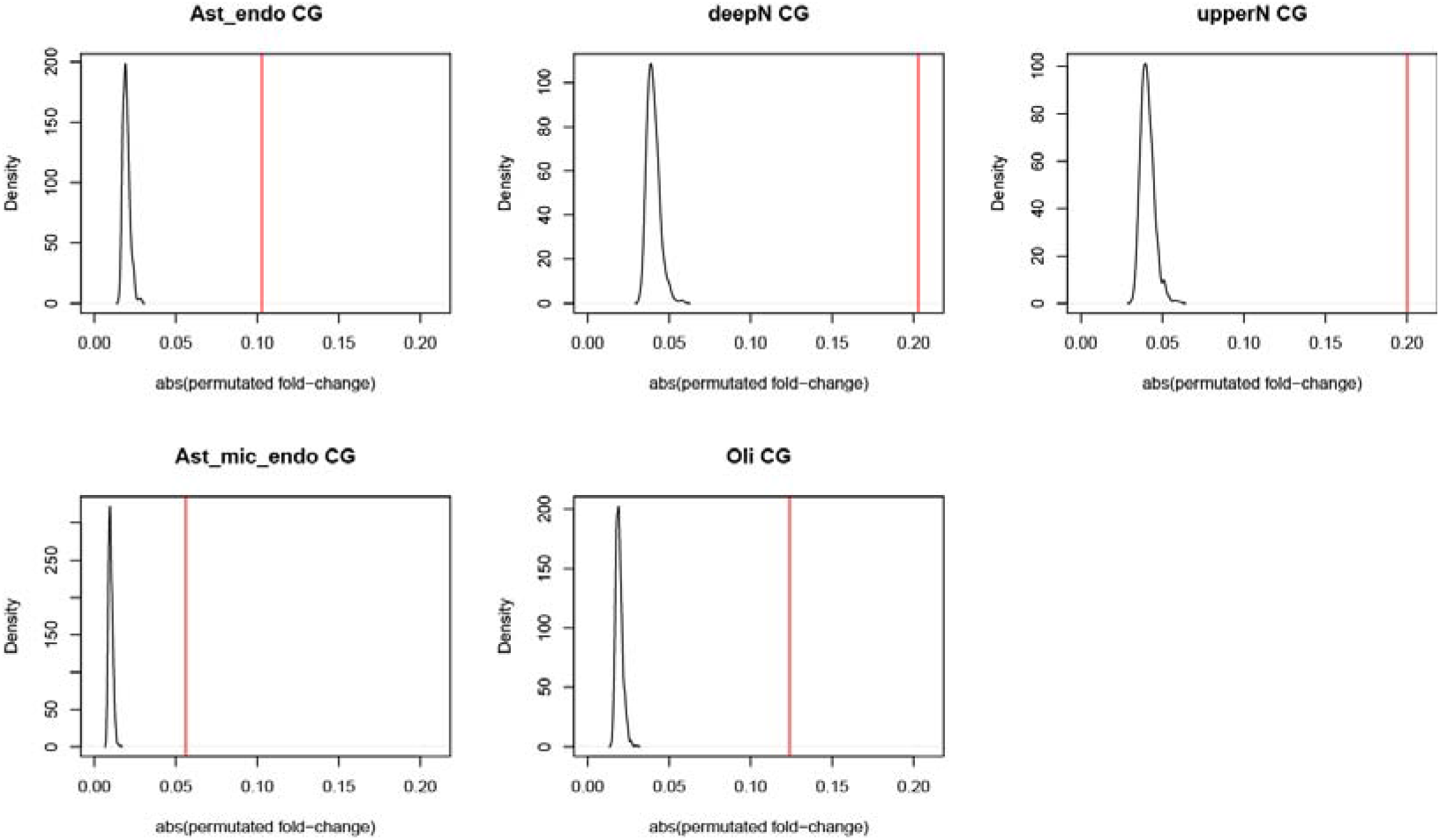
Permutation test of fold changes of differentially expressed genes in schizophrenia. The sample labels were shuffled and fold changes were calculated from two randomly selected groups. This process was repeated 1000 times. The black lines are the distribution of permutated fold changes and the red lines are the observed fold changes in each CG.

**Supplemental table 1** Collected brain cell marker genes

**Supplemental table 2** Differential expressed genes in SCZ per cell group

**Supplemental table 3** TWAS genes in SCZ per cell group

**Supplemental table 4** Coexpression-module member genes and their module membership in each cell group

**Supplemental table 5** Source data used in figures and tables

## Notes

### Competing Interest Statement

The authors have declared no competing interest.

